# Integrated Microfluidic Platform for High-Throughput Generation of Intestinal Organoids in Hydrogel Droplets

**DOI:** 10.1101/2025.06.14.659486

**Authors:** Barbora Lavickova, Hannah Theresa Kronabitter, Mar Cervera-Negueruela, Eylul Ceylan, Laura Benito Zarza, Ruben López-Sandoval, Irineja Cubela, J. Gray Camp, Jose L. Garcia-Cordero

**Author notes:** these authors contributed equally to this work (Barbora Lavickova, Hannah Theresa Kronabitter).

## Abstract

Organoid research offers valuable insights into human biology and disease, but reproducibility and scalability remain significant challenges, particularly for epithelial organoids. Here, we present an integrated microfluidic platform that addresses these limitations by enabling high-throughput generation of uniform hydrogel microparticles embedded with intestinal stem cells. Our platform includes a cell distribution system for homogenous cell encapsulation and a microfluidic oil removal module for efficient particle transfer to aqueous media. We demonstrate the successful culture and differentiation of both healthy and tumor-derived intestinal organoids within these microparticles, achieving high homogeneity and reproducibility. This integrated microfluidic approach holds promise for scalable and standardized organoid production, with potential applications in drug screening, disease modeling, and personalized medicine.

## Introduction

Organoid research stands at the forefront of biomedical innovation, providing unprecedented insights into human development and disease. However, the lack of reproducibility and scalability in organoid production, especially for epithelial organoids, such as intestinal organoids that are typically cultured in 3D-shaped hydrogel domes, presents a significant obstacle. Organoids embedded in hydrogel are often difficult to manipulate^1,2^, and 3D structure of hydrogel dome makes upstream image analysis of individual organoids challenging^3–5^. Furthermore, due to spatiotemporal gradients of growth factors and oxygen they exhibit heterogeneity in size, shape, and cell composition ^2,6–9^. Droplet-based microfluidics allows for the generation of thousands of small uniform particles, therefore offering a promising solution for high-throughput generation of organoids with both capsules and hydrogel droplets/microparticles, demonstrating significant potential for standardization of organoids and their use in drug screenings^10,11^.

While capsules have the potential to be used for suspension organoids as brain^12^ and pancreatic islet ^13^, hydrogel-based droplets/particles are often used for embedded epithelial organoids. The micrometer hydrogel particles are frequently formed using high-stiffness synthetic hydrogels, such as gelatin methacryloyl (GelMA)^14,15^ and alginate^16,17^. While these materials are user-friendly, as they can be polymerized within seconds by UV or ion bath and are not temperature-sensitive, they often inadequately support organoid growth. Alternatively, basement membrane-like matrices such as Matrigel are sometimes used^18–20^. However, these matrices present challenges due to their temperature sensitivity and insufficient droplet stability, which can hinder their long-term use and applicability in bioreactors. Moreover, many encapsulation methods struggle to ensure uniform cell distribution within droplets and capsules^19^, due to cell clamping and aggregation. This can be overcome by a high cell densities encapsulation^21,22^, addition of magnetic stirrer ^23^, or distribution device ^24^.

Although oil-free droplet generation devices are gaining popularity^13,25^, they are more complex^21,26^, and rely on specific material composition^11,13^. Therefore, oil based droplet microfluidics methods, such as flow-focusing, remain the most popular and most straightforward techniques to produce highly monodisperse droplets across a wide range of volumes^10^. Depending on the type of particle, various oil removal methods^27^ such as demulsification^28,29^, oil absorption by membranes^30,31^, and filtration^32^, are employed. These methods are manual and may result in cell death, particle loss, and oil residues. To address these issues on-chip oil removal solutions have been introduced^33^. However, these devices were developed for fast polymerization UV-based hydrogels^14^, high-viscosity oils, and high-stiffness particles, incompatible with organoid growth.

To address these challenges, we introduce a microfluidic platform featuring an integrated cell distribution system and a particle generation device, that enables the controlled production of thousands of highly uniform hydrogel microparticles embedded with primary-derived adult stem cells, serving as precursors for organoid growth. We demonstrate that optimizing hydrogel composition is essential for high-efficient cultivation of epithelial organoids, while maintaining the stability for bioreactor cultivation, manipulation, and distribution to well plates. Lastly, our microfluidic oil removal module facilitates the transition of microparticles from oil to aqueous media, minimizing manual handling and particle loss, thereby enhancing the overall efficiency and scalability of organoid production.

## Results

### Generation of homogeneous hydrogel droplets

We hypothesized that achieving uniform mature organoids requires the initial generation of homogenous hydrogel droplets with consistent volumes and even distribution of cells within each droplet. To accomplish this, we employed a flow-focusing microfluidic device (Figure 1a-b, Supplementary Figure 1a-b). This device facilitated the rapid production of thousands of highly uniform hydrogel droplets (Figure 1c) within minutes, each with a volume in the nanoliter range. The particle diameters can be controlled by varying the flow rates of the oil and the hydrogel precursor, as well as by modifying the dimensions of the microfluidic channels (data not shown). For our purposes, we chose a droplet diameter of approximately 350 µm, which is optimal for generating intestinal organoids of significant size while retaining ease of manipulation.

**Figure 1.**
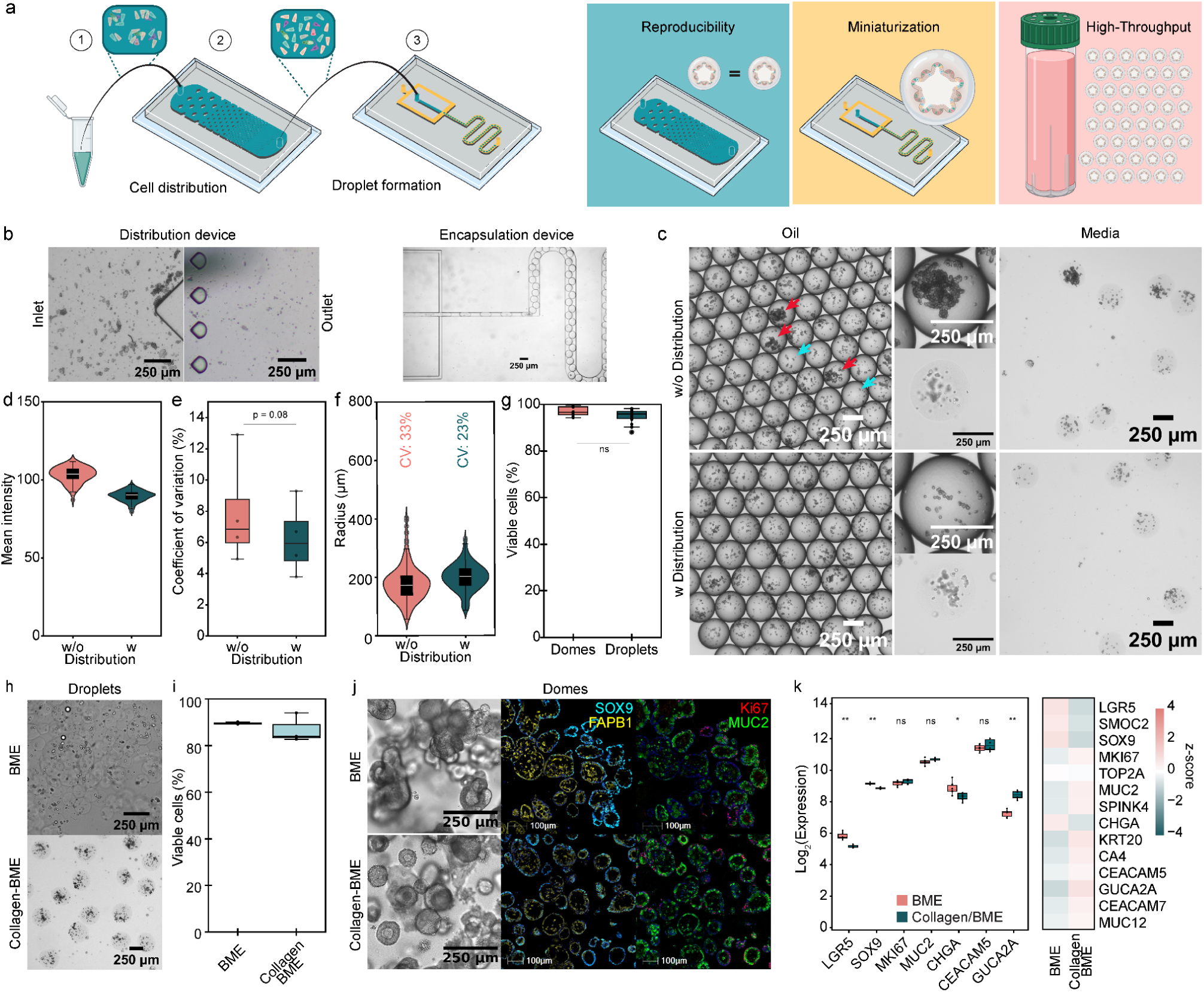
Generation of organoid droplets: a) Schematics of the microfluidic chips used for droplet generation. A single cell solution is flushed through a distribution chip to ensure cell homogeneity. Droplets are generated in a flow-focusing device where oil disrupts the hydrogel flow. This allows for high-throughput generation of miniaturized hydrogel culture systems, with increased organoid reproducibility. Schematic was partially created with BioRender.com. b) Brightfield images of cell distribution device pillars of tear-drop shape at the inlet and outlet. The cell encapsulation device with the flow focused generation of cell-hydrogel droplets (scale bars 250 µm). c) Brightfield images of droplets in oil and media, showing the cell density with and without cell distribution device (scale bars 250 µm). Droplets densely packed with cells are indicated by red arrows, nearly empty droplets are indicated with blue arrows. d) Mean intensities of droplets generated with and without the cell distribution device on day 0. Data extracted from brightfield images. e) Quantitative comparison of the coefficient of variation (CV) in cell laden droplets on day 0 across experiments. Each dot represents the coefficient of variation in a single experiment (n = 4 biological replicates). f) Size analysis of colon organoids on day 8 formed with or without distribution device, with coefficient of variation (CV) highlighted above (n_organoids_ ≥ 300). g) Quantitative fluorescence analysis of cell viability of cells after encapsulation on days 0 in domes or droplets by live/dead staining (n_samples_ = 4). h) Brightfield images of cell-hydrogel droplets composed of 100% BME (Matrigel) and 1.5 mg/mL Collagen I - 25% BME (Matrigel) ECM mix (scale bars 250 µm). i) Quantitative fluorescence analysis of cell viability of intestinal organoids at days 7 of culture in BME (Matrigel) and 1.5 mg/mL Collagen I / 25% BME (Matrigel) ECM mix domes by live/dead staining (n_samples_ = 3). j) Brightfield images of healthy colon organoids grown on day 6 (scale bars 250 µm) and mIF staining of healthy colon organoids grown on day 8 (SOX9 turquoise, FABP1 yellow, MUC2 green and Ki67 red, scale bars 100 µm) comparing the two hydrogels, BME (Matrigel) and 1.5 mg/mL Collagen I - 25% BME (Matrigel) ECM mix. k) Quantification of Log2 transformed expression levels and heatmap of Z-score normalized expression levels across selected genes for the two ECM conditions. Data were averaged across replicates. Statistical significance between conditions was assessed using the Wilcoxon test, with p-values adjusted for multiple testing using the Benjamini-Hochberg method. P-value of < 0.05 was expressed as *, p-value < 0.01 was expressed as ** and p-value < 0.001 was expressed as ***. The normality of the distribution was verified with a Shapiro-Wilk test and a t-test was done to calculate the p-values.

While microfluidic droplet formation resulted in highly homogeneous hydrogel droplets, we observed variability in the number of encapsulated cells within the droplets, with some droplets being densely packed with cells while others were nearly empty (Figure 1c). High-speed camera observations revealed a pulsatile flow caused by cell clumping and inhomogeneous cell dispersion within the hydrogel precursor. To address this issue, we implemented a microfluidic distribution device featuring a series of pillar arrays with varying diameters and pitches prior to encapsulation. This design, inspired by a previously reported device^34^, aimed to break up cell clumps and homogenize cell distribution in the precursor mixture (Figure 1a). However, we observed that the circular shape of the pillars led to significant device clogging (data not shown). To mitigate this issue, we optimized the pillar design by adjusting their shape, size, and spacing (Supplementary Figure 1c). Tear-drop-shaped pillars proved to be the most effective, as they maximized cell yield and minimized clump formation. At the entrance of the distribution device, larger cell clumps were evident (Figure 1b), but at the exit, these clumps were significantly reduced (Figure 1b). This optimization resulted in hydrogel droplets with a more uniform cell distribution and no visible high-density clumps (Figure 1c).

The improved cell distribution was confirmed by measuring the mean pixel intensity in droplets, which serves as an approximation of cell densities. On day 0, droplets produced using the distribution device showed reduced dispersion and greater consistency in mean pixel intensity compared to the one generated without the device (Figure 1d). This was further supported by a reduced coefficient of variation in the mean intensities of droplets generated with the distribution device across different experiments (n=4) (Figure 1e). By day 8, organoids in droplets generated with the distribution device exhibited a 10% lower coefficient of variation compared to those generated without it (Figure 1f), confirming increased reproducibility and reduced size variation. Additionally, high cell viability (95-100%) immediately after encapsulation, with no significant difference compared to cells directly seeded in dome culture (Figure 1g), indicates that the encapsulation process imposes low stress on the cells.

Basement membrane-like matrix (BME) as Matrigel and Cultrex is the most commonly used matrix for intestinal organoid growth and has been previously used for hydrogel droplet formation^19–22^. However, its soft properties result in unstable droplets (Figure 1h), complicating oil extraction and leading to droplet merging and disintegration over time. To enhance droplet stability while maintaining support for epithelial organoid culture ^35^, we added type I collagen to the BME. We tested different hydrogel compositions by adjusting the collagen-to-BME ratio (Supplementary Figure 2). Live/dead staining of intestinal organoids grown in different hydrogel mixtures on day 7 (Figure 1i, Supplementary Figure 2a-b) indicated that the highest viability, comparable to pure BME, was achieved with a mixture containing 1.5 mg/mL collagen and 25% BME, resulting in 85% viability. Comparable viability was also observed with a 50% BME concentration. However, higher collagen concentrations (2.5 mg/mL) resulted in lower cell viability (60-70%). We selected a mixture of 1.5 mg/mL collagen supplemented with 25% BME, which maintained high cell viability and produced stable droplets with higher integrity (Figure 1h), preventing droplet merging, simplifying oil removal, and allowing for centrifugation and easy manipulation. Additionally, the droplets remained stable over time, supporting both static and bioreactor cultures. Organoids grown in this extracellular matrix (ECM) mixture displayed a dense morphology with elongated columnar cells characteristic of cell differentiation in intestinal organoids, with less budding morphology observed compared to those grown in pure BME (Figure 1j). Multiplex immunofluorescence (mIF) staining (Figure 1j) confirmed the presence of diverse cell types, including stem cell and proliferation markers (SOX9^+^, Ki67^+^), enterocytes (FABP1^+^), and goblet cells (MUC2^+^). Bulk RNA sequencing (Figure 1k, Supplementary Figure 2c) of organoids grown in ECM mix or BME revealed similar expression patterns, with slight downregulation of stemness markers (LGR5, SOX9) and upregulation of FABP2, GUCA2A, indicating a slightly higher level of differentiation in the ECM mix domes.

### Organoid development and homogeneity

To assess the biocompatibility of the encapsulation process and determine if the organoids resembled those grown in traditional culture and their parental tissues, we cultured droplets containing colon or duodenum intestinal adult stem cells in a bioreactor for 8 days, and performed brightfield imaging, Hematoxylin and Eosin (H&E), and mIF staining to evaluate their development. By day 3, the single cells formed aggregated organoids (r = 75 µm) that began to grow and differentiate. By day 6, these organoids grew in size (lumen r = 118 µm) and exhibited a proliferative phenotype marked by SOX9 and Ki67 expression. By day 8, the organoids further increased in size (r = 160 µm) and matured as indicated by the presence of strong FABP1 expression (Figure 2a-b). Similar results were observed for proximal small intestinal organoids from the duodenum (Supplementary Figure 4a-b). Organoids cultured in the bioreactor remained individual and mostly uniform through day 8. In contrast, static organoids grown in a 6-well plate merged over time, forming larger, more heterogeneous structures (Supplementary Figure 3). The whole-mount staining of F-actin and collagen on day 8 (Figure 2c) demonstrates individual organoid development supported by the hydrogel particles. Furthermore, the size of the organoids could be modulated by varying the numbers of cells per droplet (Figure 2d). Specifically, 5000 cells/µL (approximately 130 cells/droplet) resulted in organoids with a radius of ∼120 µm, while 7500 cells/µL (approximately 200 cells/droplet) produced larger organoids with a radius of ∼230 µm. A key benefit and hallmark of the intestinal organoid model is the ability to differentiate into various intestinal epithelial cell types. mIF of day 8 organoids (Figure 2e, Supplementary Figure 4c) confirmed the presence of diverse cell types, including stem cells (SOX9^+^), transit-amplifying cells (Ki67^+^), colonocytes (FABP1^+^, CA1^+^) or enterocytes (FABP1^+^, APOA4^+^), enteroendocrine cells (CHGA^+^), and goblet cells (MUC2^+^). We observed a higher number of goblet cells (MUC2^+^) in colon organoids and a higher prevalence of enteroendocrine cells (CHGA^+^) in small intestine organoids, which is concordant with previous reports^36,37^. qPCR (Figure 2f) and image analysis of mIF staining (Figure 2g, Supplementary Figure 4d) of the main canonical markers over time revealed downregulation in the expression of stem cell markers (LGR5, SOX9), stable expression of proliferation markers (MKI67/Ki67), and upregulation of differentiation markers for goblet cells (MUC2^+^), enteroendocrine cells (CHGA^+^), and enterocytes / colonocytes (FABP1^+^; FABP1^+^, CA1^+^) by day 8, which highlight the ability of the organoids to homeostatically grow and differentiate within the droplets.

**Figure 2.**
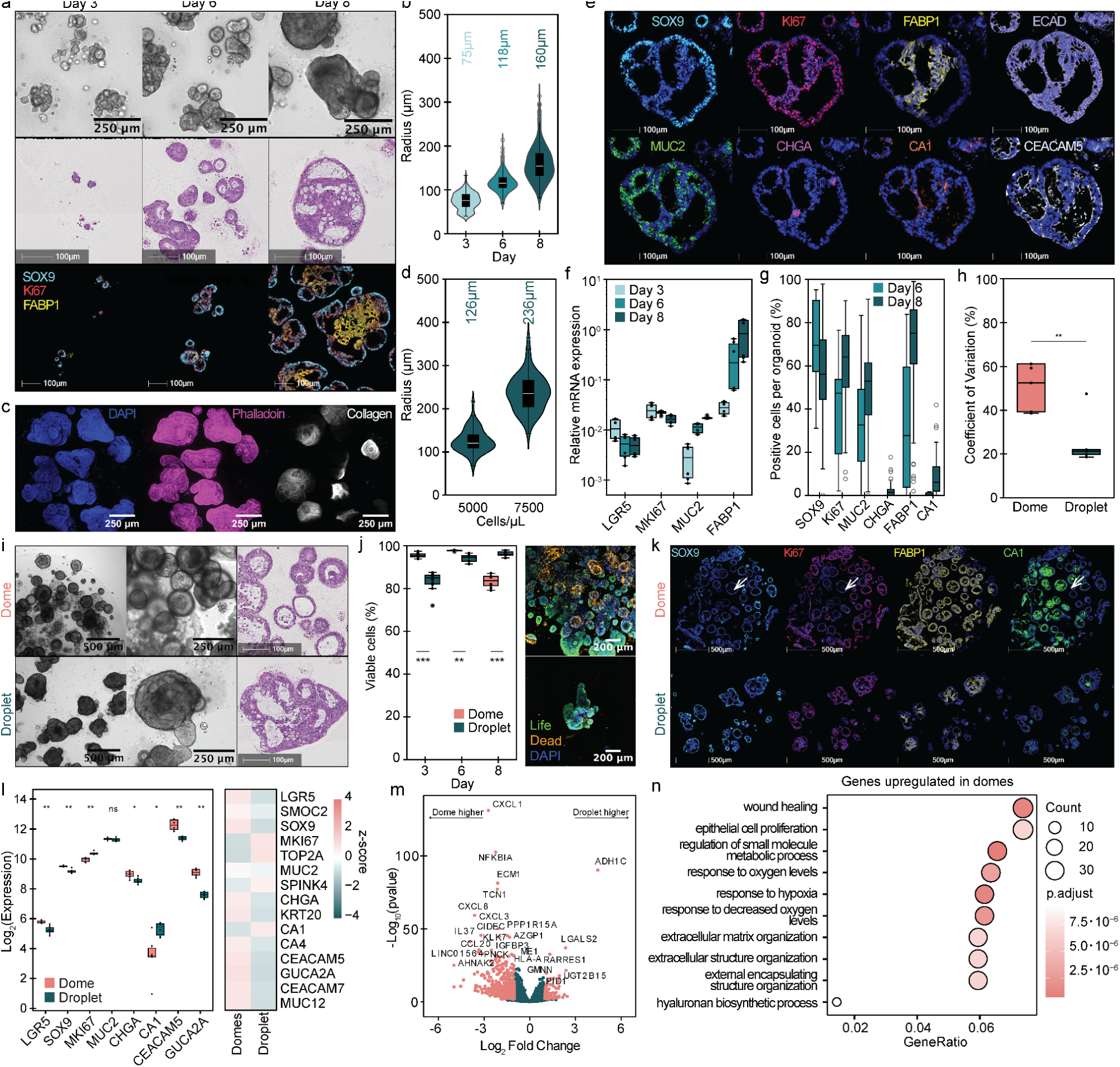
Development of colon organoids in droplets and comparison to traditional dome cultures: a) Brightfield images (scale bars 250 µm), H&E staining (scale bars 100 µm), and mIF staining (SOX9 turquoise, FABP1 yellow, Ki67 red, scale bars 100 µm) of colon organoids in hydrogel droplets on days 3, 6, and 8. b) Size development analysis of colon organoids over several days, with average size highlighted above (n_organoids_ ≥ 235). c) Whole-mount staining of colon organoids in droplets on day 8. d) Size development analysis of duodenum organoids formed from different amounts of encapsulated cells, with average size highlighted above (n_organoids_ ≥ 58). e) mIF staining of organoids in hydrogel droplets on day 8 (SOX9 turquoise, FABP1 yellow, Ki67 red, MUC2 green, CA1 orange, CHGA pink, ECAD purple, CEACAM5 white, scale bars 100 µm). f) Relative mRNA expression of different genes of organoids grown in droplets over several days normalized to β-actin gene expression (2 biological replicates). g) Quantification of mIF staining of organoids in droplets on day 6 and day 8 (n_organoids_ ≥ 30). h) Quantitative comparison of the coefficient of variation in organoid sizes between organoids grown in domes (1:12 dilution) and droplets on day 6. Each dot represents the coefficient of variation in a single experiment (n = 5 biological replicates). i) Brightfield images (scale bars 500 and 250 µm) and H&E staining (scale bars 100 µm) of organoids grown in domes and droplets on day 8. j) Quantitative fluorescence analysis of cell viability in intestinal organoids at days 3, 6, and 8 of culture in domes or droplets by live/dead staining (n_samples_ = 4), and representative fluorescent images of organoids grown in domes and droplets on day 8. k) mIF staining of organoids in domes (1:12 dilution) and hydrogel droplets on day 8 (SOX9 turquoise, FABP1 yellow, Ki67 red, CA1 green, scale bars 100 µm). l–n) Bulk RNA sequencing of organoids grown in domes (1:6 dilution) and droplets (3 technical replicates, 2 biological replicates). l) Quantification of Log2 transformed expression levels and heatmap of Z-score normalized expression levels across of selected genes. Data were averaged across replicates. Statistical significance between conditions was assessed using the Wilcoxon test, with p-values adjusted for multiple testing using the Benjamini-Hochberg method. m) Volcano plot of differentially expressed genes between domes (left) and droplets (right) subsets (p < 0.05, log2 Fold Change > 1). n) Plot showing significantly enriched Gene Ontology biological processes of genes in colon organoids grown in domes (p-value < 0.05, q-value < 0.2). P-value of < 0.05 was expressed as *, p-value < 0.01 was expressed as ** and p-value < 0.001 was expressed as ***. The normality of the distribution was verified with a Shapiro-Wilk test and a t-test was done to calculate the p-values.

To further characterize the organoids and benchmark them to the traditional dome culture, we compared cells encapsulated in hydrogel droplets to those cultured in domes (Figure 2h-i). However, due to differences in volume and shape, monitoring and segmenting individual organoids in domes is challenging^3^, especially in densely populated cultures. Additionally, organoids can be located in various z-planes, making segmentation difficult due to image overlapping structures at various heights. This necessitates significant dilution of the cell suspension prior to dome seeding to prevent organoid merging and allow for individual monitoring, which in turn requires large and often costly volumes of hydrogels and media. While we diluted the cell suspension for dome culture 6- to 12-fold, this still often resulted in difficulties in organoid segmentation at later time points (Supplementary Figure 5). Therefore, in this study, we compared organoid sizes on day 6. Organoid generation in droplets showed high reproducibility across biological replicates (Supplementary Figure 6a), with an average coefficient of variation (CV) of 20% for droplet-derived organoids compared to 50% for dome-grown organoids (Figure 2h). Organoids formed in droplets therefore grow significantly larger (r_dome_ = 46 µm, r_droplet_ = 118 µm) and more uniformly (CV_dome_ = 59%, CV_droplet_ = 19%) (Figure 2i, Supplementary Figure 6b).

In intestinal organoid culture, larger domes create steep nutrient, growth factors and oxygen gradients, leading to poorly perfused regions and resulting in variability in organoid size, morphology, and development ^2,6–9^. Although various dome volumes are utilized in the literature, larger domes are prone to developing steep gradients. To enable meaningful comparison between organoids grown in droplets and domes, we hypothesized that smaller domes would reduce these gradients and produce more uniform organoids. Therefore, we limited our dome cultures to 3.5 µL, a volume typically lower than that used in standard dome cultures, in order to achieve sufficient dome stability while minimizing these gradients. Nevertheless, we were able to observe differences between both culture approaches in terms of homogeneity, viability and cell type localization. Although initial cell viability was lower in droplet cultures on day 3, likely due to stress from encapsulation, this pattern reversed by day 8 (Figure 2j, Supplementary Figure 6c). The increased viability in droplets compared to domes by day 8 is likely due to higher proliferation rates and improved nutrient supply in the bioreactor droplet culture system. Live/dead fluorescence images showed a high number of dead cells at the center and increasing viability towards the edges (Figure 2j), indicating the presence of a necrotic core within the dome. Similarly, mIF stainings (Figure 2k) revealed notable differences in marker distribution between organoids grown in domes and those in droplets. In dome culture, the edges exhibited high SOX9 and Ki67 expression, indicating proliferation, whereas the center showed elevated levels of FABP1 and CA1 expression, suggesting increased differentiation due to limited growth factors. This pattern is well-documented^2,6,9^ and reproducible. In contrast, organoids in droplets contained markers for both proliferation and differentiation, resulting in more homogeneous organoids. While significant differences were observed in individual patterns, mIF analysis (Supplementary Figure 6d) showed less variability between groups overall. Organoids grown in droplets displayed slightly lower expression of differentiation markers (FABP1, CA1). To further investigate these differences, we performed bulk RNA sequencing (RNA-seq) of day 8 organoids. Both groups displayed similar profiles of standard colon-specific marker genes (Figure 2l), with differentiation markers slightly lower, and the proliferation marker (MKI67) upregulated in the droplet-grown organoids, consistent with mIF and live/dead analysis. Interestingly, differential analysis (Figure 2m) revealed the presence of cytokine and chemokine genes (CXCL1, CXCL8, CCL20) in dome conditions. Gene Ontology analysis of upregulated genes in domes (Figure 2n) indicated enrichment of genes associated with wound healing, hypoxia response, and extracellular matrix reorganization, aligning with the lack of oxygen and nutrient supply in dome centers. We also observed upregulation of genes associated with hypoxia response in droplets cultured in static conditions (Supplementary Figure 6e-g), highlighting the importance of enhanced nutrient and gas exchange within the bioreactor.

Altogether, these findings confirm the biocompatibility and the feasibility of 3D cell culture using hydrogel scaffolds, and underscore the advantages of culturing epithelial organoids individually in small droplets, to provide a more controlled microenvironment leading to higher reproducibility and homogeneity. Moreover, integrating this method with a bioreactor setup offers potential for high-throughput organoid generation, paving the way for scalable and standardized organoid production.

### Tumor organoids development in hydrogel droplets

Having established droplet bioreactor culture for healthy intestinal organoids, we further applied the technique to patient-derived tumor organoids (Figure 3a). Similar to healthy organoids, by day 3, cell aggregates with an average radius of 71 µm had formed in the hydrogel droplets, which further developed into 3D structures with an average radius of 119 µm over the initial 6 days. Further tumoroid cultures did not lead to an increase in the size of the organoids (Figure 3a, b), but development of dense aggregates. H&E staining of the tumoroids confirmed their dense structure with a lack of luminal space (Figure 3c). This aligns with the fact that patient-derived tumoroid cultures are known to adopt donor-specific morphologies^38^. As expected, tumoroids grew larger in droplets due to the larger cell concentration, while tumoroid sizes in domes were significantly smaller (Figure 3c). In contrast to healthy intestinal organoids, which showed clear patterning and differentiation over time, tumoroids did not show a patterned cell distribution (Figure 3d). This is likely due to the lower amount of growth factors in the tumor organoid media, and particularly the lack of Wnt, which has been linked to the observed patterning^9^. To evaluate the potential use of the generated tumoroids as a screening platform, we transferred the fully developed organoid droplets to a 96-well plate and treated them with SN-38^2^, chemotherapy drugs Capecitabine, combination chemotherapy drug FOLFOX (5-Fluorouracil:Leucovorin:Oxaliplatin 25:5:1)^21,39^ and target therapy drug Bortezomib^40^ (Figure 3e). The tumoroids showed sensitivity to SN-38 and Bortezomib, reflected by an induced cytotoxic effect with increasing concentrations. In contrast, we did not observe cytotoxicity changes when using FOLFOX and Capecitabine. Overall, these results demonstrate that the system can be adapted to different organoid cultures and highlight the potential use of the platform for chemotherapy screening applications.

**Figure 3.**
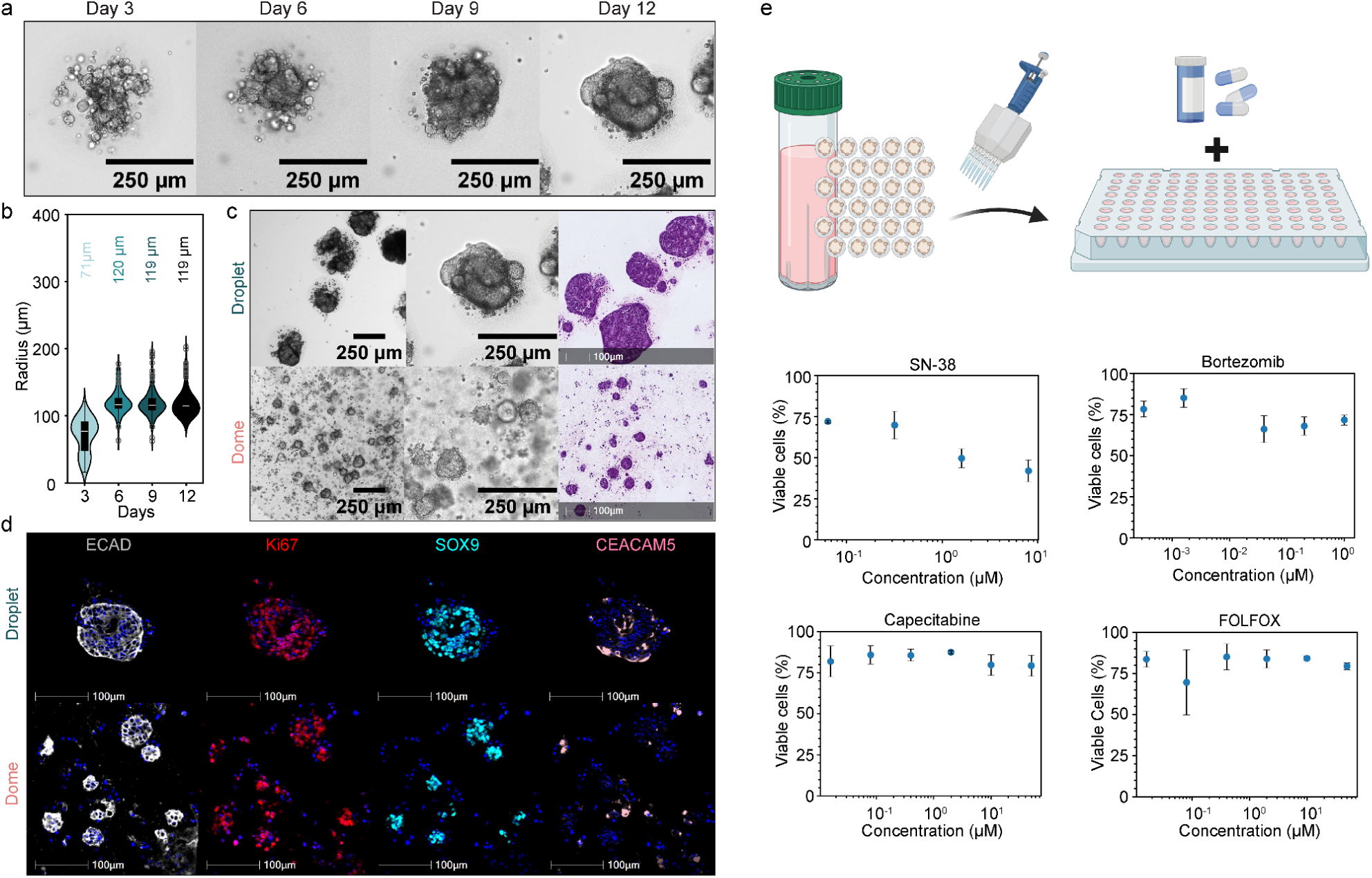
Growth and morphology of colon tumor organoids: a) Brightfield images of colon tumor organoids in hydrogel droplets on day 3, 6, 8 and 12 (scale bar 250 µm). b) Size development analysis of colon tumor organoids over several days, with average size highlighted above (n_organoids_ < 110). c) Brightfield images (scale bar 250 µm) and H&E staining (scale bar 100 µm) of organoids grown in domes (1:6 dilution) or droplets. d) mIF staining of organoids in hydrogel droplets and domes (1:6 dilution) on day 12 (ECAD white, Ki67 red, SOX9 turquoise, CEACAM5 pink, scale bars 100 µm). e) Drug treatment of bioreactor grown tumor organoids. Dose-response curves to compounds FOLFOX (5-Fluorouracil,: Leucovorin:Oxaliplatin = 25:5:1, the concentration specified in the graph is based on 5-Fluorouracil), SN-38, Bortezomib, Capecitabine, quantified by fluorescence analysis of cell viability by live/dead staining (n_samples_ = 3). Schematic was created with BioRender.com.

### Automated oil extraction device and organoid spotting

Although microfluidic droplet generation is a well-characterized and automated process, the subsequent steps of particle polymerization and oil removal are often performed manually^27^. Traditional oil removal methods, such as oil absorption by hydrophobic membranes^30,31^ or using demulsifiers to chemically break emulsions^28,29^, involve significant manual handling. These processes are highly user-dependent and can lead to material loss. To address these challenges and to further automate the organoid generation process, we integrated the system with a polymerization chamber and a microfluidic oil removal device adapted from Mohamed et. all^14^. The modifications allowed for a handling of temperature-sensitive soft hydrogel particles and low surface tension oils (Figure 4a, Supplementary Figure 7a). After droplet generation, the aqueous precursor droplets were introduced into the polymerization chamber (Supplementary Figure 7b), where they were maintained at 37°C for 20 minutes. This chamber featured two outlets: one for waste disposal and another one connected to the oil removal device that leverages particle buoyancy. During droplet generation, the bottom waste outlet remained open, allowing oil to escape while droplets floated on top of the oil. Following polymerization, the waste outlet was closed, and the chip inlet was opened to direct the flow of polymerized particles into the oil removal device. The oil containing hydrogel particles was co-introduced into the oil removal device (Supplementary Figure 7c) along with oil containing the perfluorooctanol (PFO) demulsifier, delivered using pulse-width flow modulation. The demulsifier and oil were then mixed within a serpentine channel, where PFO disrupted the surfactants at the oil-water interfaces of the droplets, destabilizing them. Consequently, when particles entered the extraction oil-media coflow chamber, they exhibited a higher affinity for the aqueous medium, transitioning from the oil phase to the aqueous solution for collection.

**Figure 4.**
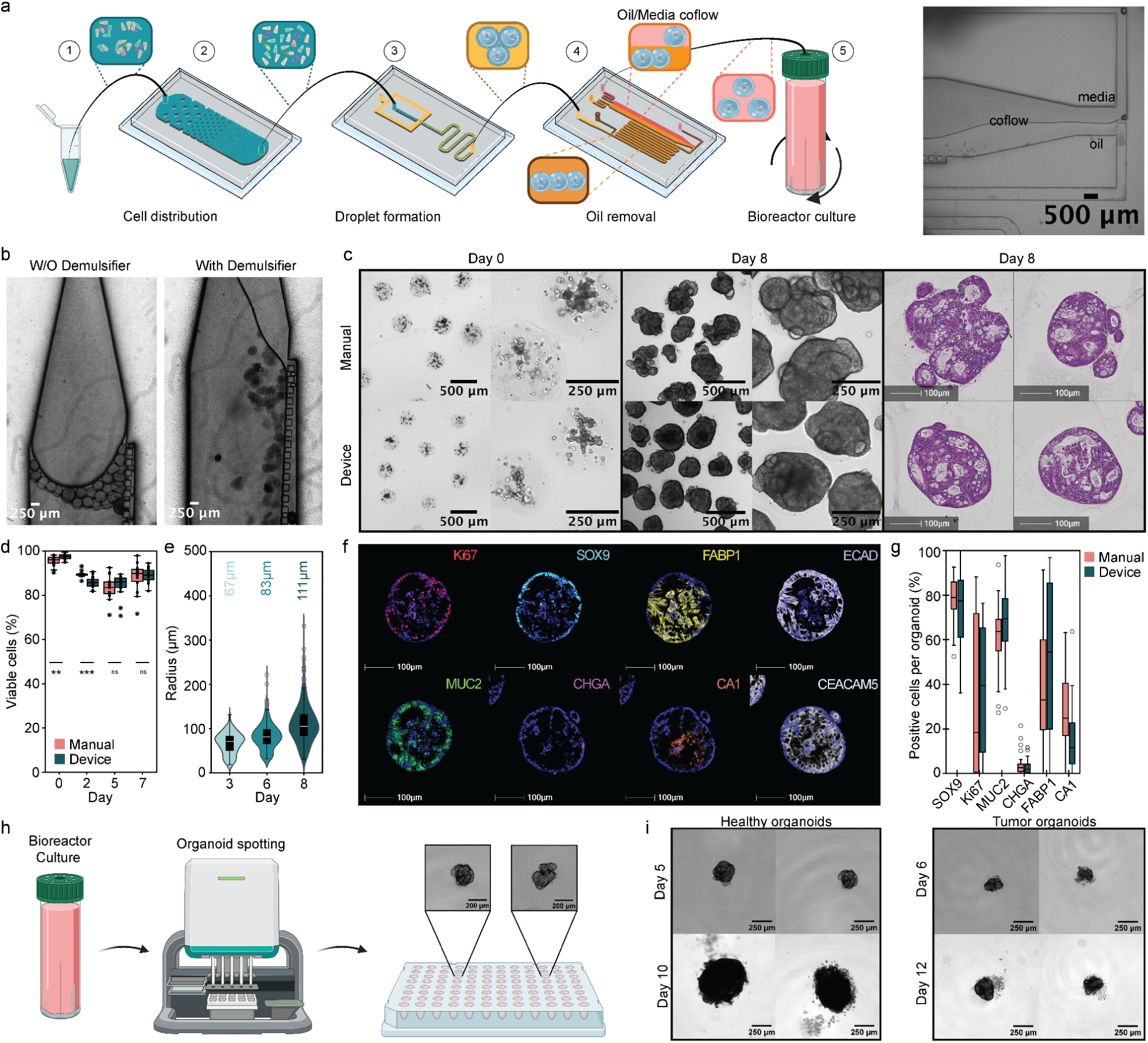
Oil Removal Device and Organoid Spotting: a) Schematic overview of oil removal device integration. Schematic was partially created with BioRender.com. Brightfield image of coflow of oil and water fraction in extraction chamber (scale bars 500 µm). b) Brightfield images of droplets in extraction with or without demulsifier pulsed into the device (scale bars 250 µm). c) Comparison of hydrogel droplets generated with manual oil removal method or with the oil removal device. Brightfield images of encapsulated cells on day 0 and brightfield images (scale bars 500 µm and 250 µm) and H&E staining (scale bars 250 µm) of colon organoids in hydrogel droplets on day 8. d) Quantitative fluorescence analysis of cell viability of intestinal organoids generated by manual oil removal or with oil removal device at days 0, 2, 5, and 7 of culture by live/dead staining (n_organoids_ =11). d) Size development analysis of organoids generated with the oil removal device over several days, with average size highlighted above (n_organoids_ ≥ 240). f) mIF staining of organoids in hydrogel droplets generated with the oil removal device on day 8. g) Quantification of mIF staining of organoids in droplets with manual oil removal method or with oil removal device (n_organoids_ ≥ 45). P-value of < 0.05 was expressed as *, p-value < 0.01 was expressed as ** and p-value < 0.001 was expressed as ***. The normality of the distribution was verified with a Shapiro-Wilk test and a t-test was done to calculate the p-values. h) Schematic of organoid distribution from bioreactor to a well plate by organoid spotting. i) Representative brightfield images of organoids and tumoroids development after distribution in a well plate (scale bars 250 µm).

The extraction process relied heavily on the presence of a demulsifier (Figure 4b), without it, droplets retained their oil affinity, leading to pulsatile flow or droplets being squeezed through pillars towards the oil outlets. However, in the presence of the demulsifier, droplets transitioned smoothly to the media or buffer for collection (Figure 4b). The device’s geometries (Supplementary Figure 7d) were optimized for maximal droplet yield, minimal clogging, and reduced oil carryover. Key factors for effective coflow establishment and particle transfer included a higher chamber-to-input channel width ratio (1:6), a low oil-to-media flow ratio, and a tapered shape at the chamber entrance, which also helped minimize particle attachment to the chamber or microposts. Overall, the device exhibited low oil transfer and reduced droplet loss compared to manual methods. Despite being composed of a soft hydrogel, the droplets maintained their integrity post-oil extraction (Figure 4c). Furthermore, cell viability following the extraction process was similar to manual extraction, exceeding 95% on day 0, suggesting that the device does not induce additional stress on the cells and that the incubation time with PFO has minimal cytotoxic effects on the resulting particles (Figure 4d). As anticipated, the encapsulated cells developed into colon organoids with diameters <200 µm over eight days in culture (Figure 4e), exhibiting morphology similar to those extracted manually or grown in standard dome culture (Figure 4c, Supplementary Figure 8). Additionally, mIF staining of the droplets revealed characteristic colon organoid markers (Figure 4f-g), consistent with those obtained via manual methods (Figure 2c). These results indicate that the integrated microfluidic approach is readily applicable for generating hydrogel droplets suitable for single organoid growth.

Lastly, to simplify the workflow, we tested if our microfluidic-based pipeline can work alongside liquid handling tools to transfer suspended organoids to multiple well plates. This facilitates a seamless process from organoid generation to assay, allowing for applications such as drug screening, organoid imaging, and analysis typically performed in well-plate formats. Thanks to the stability of the hydrogel particles, this transfer can happen either immediately after organoid precursor generation or later during culture. The latter method offers advantages, including improved oxygenation and nutrient supply in the bioreactor, easier media exchange, and the potential for organoid sorting prior to distribution. To assess the feasibility of transferring individual organoids, we used a commercially available benchtop dispenser robot (Dispen3D, Seed Bioscience, Figure 4h) to transfer organoids from the bioreactor to 96 well-plates. Our results showed that the dispensing process does not compromise organoid viability, as they continue to grow after being transferred to the well plate (Figure 4i). This capability opens up future possibilities for applications such as drug testing, or co-culture with individual or specific numbers of organoids.

In conclusion, the stability of the ECM mixture and the microparticles allows for a combination of different automation methods and high throughput droplet generation. These automation methods minimize manual handling and simplify organoid manipulation, which has potential to further enhance reproducibility of organoids and expand their applications.

### Conclusion

Here, we introduced an integrated microfluidic process for generating hydrogel droplets that enables 3D culture, growth, and large-scale production of intestinal organoids and tumoroids. By incorporating a dissociation device before the droplet formation unit, we achieved more uniform cell distribution within the droplets. The integration of a hydrogel polymerization chamber and an oil removal device after droplet generation allows for hands-free hydrogel production, eliminating manual handling and reducing droplet loss. Additionally, we optimized the ECM composition of the droplets to ensure their stability over time while maintaining proper organoid development, allowing for the cultivation of organoids in bioreactors and facilitating easy handling for downstream applications, such as transferring to well plates.

The established system reduces variability in organoid culture by providing uniform 3D scaffolds in a reproducible manner. Organoids grown in these droplets showed higher individual homogeneity in size and distribution of canonical markers compared to traditional dome cultures, which often exhibit higher differentiation and cell death in the center, and higher stemness at the edges of the dome. Furthermore, we observed lower expression of cytokines and genes associated with hypoxia response in droplet cultures, highlighting the importance of enhanced nutrient and gas exchange within the droplets. These findings underscore the advantages of culturing epithelial organoids individually in small droplets, creating a more controlled microenvironment that enhances reproducibility and homogeneity.

We believe this work provides proof-of-concept for an integrated microfluidic process for generating hydrogel droplets for human organoid and tumoroid culture. This system has the potential to be applied to various epithelial cell organoids, as well as co-culture systems involving stromal cells or immune cells. Our results suggest that our approach opens up potential avenues for scalable, high-throughput, and reproducible culture of diverse epithelial organoids and tumoroids. Such capabilities are particularly valuable for advancing application potential in drug screening, genetic engineering, and personalized medicine, where consistent and efficient organoid production is essential.

## Materials and Methods

### 1. Material

All the material is specified in Supplementary Table 1.

### 2. Human tissue samples

List of donors is specified in Supplementary Table 2. Human intestinal tissue resections, along with concurrent data collection and experimental procedures, were conducted within the framework of the non-profit foundation HTCR (Munich, Germany) and University Center for Gastrointestinal and Liver Disease (Clarunis; Basel, Switzerland), including informed patient consent. The framework of the HTCR Foundation has been approved by the ethics commission of the Faculty of Medicine in the Ludwig Maximilian University (no. 025-12) and the Bavarian State Medical Association (no. 11142). The framework of the University Center for Gastrointestinal and Liver Disease was approved in accordance with the Helsinki Declaration and reviewed and approved by the ethics committee (Ethics Committee of Basel, EKBB, no. 2019-02118). Visceral surgery with partial resection of intestine was performed on these oncology patients, from which we used micro- and macroscopically tumor-free tissue as well as tumor material.

### 3. Organoid culture & Tumoroid culture

Human intestinal organoid lines were maintained in organoid media as previously described^41^ up until passage 20 at 37°C, 5% CO_2_ and 95% humidity incubators, with a few modifications (Supplementary Table 3). The organoids lines were regularly tested for mycoplasma by MycoAlert™ PLUS Mycoplasma Detection Kit (Lonza) according to the manufacturer’s instructions.

Tumor organoid lines were maintained until passage 26 in IntestiCult™ Organoid Growth Medium (Human, StemCell Technologies) without supplement and Advanced DMEM:F12, 1x Glutamax in a ratio of 1:1 supplemented with 0.05 mg/mL Primocin (Invivogen).

Healthy organoids and tumor organoids were fragmented by Gentle Cell Dissociation reagent (Stemcell Technologies) weekly at 1:4 - 1:8 depending on the confluence and growth rate. Washing steps were performed with Advanced DMEM:F12 (Gibco) with 0.01 M Hepes (Gibco), 1x Glutamax (Gibco) and 100 U Penicillin and Streptomycin (Gibco) referred to as DMEM:F12+++. The fragments were seeded on Tissue Culture treated well plates (CytoOne®) in 25 µL domes in 100% growth factor reduced Matrigel for healthy organoids (Corning) or Cultrex Basement Membrane Extract, PathClear for tumor organoids (R&D Systems), respectively. After 15-30 minutes polymerization in the incubator, organoid media supplemented with 1 µM Y-27632 was added. Media exchange was performed every two to three days until use.

### 4. Fabrication of microfluidic devices

The molds were generated using standard photolithography. The silicon wafers were primed in an oxygen plasma machine for 7 minutes (ATTO RF, Diener electronic GmbH), and SU-8 photoresist (GM1075, Gersteltec Sàrl) was spin-coated onto the wafer to yield the desired height. After relaxation and soft bake, the spin-coating process was repeated for wafers that required heights above 200 µm. The final heights for the different devices were as follows: distribution device (200 µm), droplet device (250 µm) (Supplementary Figure 1c), and oil removal device (400 µm, Supplementary Figure 7d). Wafers were exposed with a laser writer (µMLA, Heidelberg Instruments), followed by a post-exposure bake. The unexposed photoresist was developed using propylene glycol monomethyl ether acetate (Sigma Aldrich), and the resulting wafers were hard baked.

The microfluidic devices were fabricated with polydimethylsiloxane (PDMS, Sylgard) by soft lithography. Each of the wafers was treated with chlorotrimethylsilane (Sigma-Aldrich) and covered with a PDMS solution of 10:1 elastomer to crosslinker ratio. The molds were then degassed in a desiccator and baked at 70 °C for a minimum of 2 hours. The polymerized PDMS chips were removed from the molds, punched to create inlets and outlets, and plasma bonded to a PDMS-coated glass slide.

### 5. Organoid culture in hydrogel droplets

Organoids were washed with DMEM/F12+++, collected and centrifuged. Then, organoids were resuspended in TrypLE™ Express Enzyme (1X) (Gibco) with 350 U DNAse (Roche), 2 µM Y-27632 (STEMCELL Technologies) and 1.5-2 mM DL-Dithiothreitol solution (DTT) (Sigma-Aldrich) for 8 minutes at 37°C and an extra 5 minutes after harsh pipetting. After pelleting, cells were resuspended in DMEM:F12+++, strained through a 40 µm cell strainer (Pluriselect), centrifuged and resuspended in organoid media with 2 µM Y-27632 and counted with the Cell Countess (Invitrogen). Collagen I solution (Biomatrix, 6.2 mg/mL) was neutralized with Neutralization Solution (Biomatrix). The final extracellular matrix (ECM) solution consisted of 1.5 mg/mL neutralized collagen and 25% growth factor reduced Matrigel (Corning). Cells (5x10^6^/mL unless indicated otherwise) in organoid media were directly incorporated in the ECM mix. Then, the ECM-cell mixture was subjected to encapsulation and an oil removal process.

Organoid media was supplemented with 1 µM Y-27632 for the first three days. Droplets were cultured using the CERO bioreactor (80 RPM with 2 second rotation period). Static droplets were cultured in ultra low attachment 6 well plates (Corning). Every three days, 70% of organoid media was changed by centrifuging the droplets in media at 190 RCF for 3 minutes, prior to the media exchange.

Domes were used as controls. Therefore, we diluted the initial cell-hydrogel suspension 1:6, 1:12 and in specific cases also 1:24 and seeded two 3.5 µL domes in a well of a Tissue culture treated 24 well plate (CytoOne). The domes were polymerized for 20 minutes at 37°C and 500 µL of organoid media with 1 µM Y-27632 for the first three days was added. Every three days 100% of the media was renewed. Brightfield images were taken with the Leica DMi (Leica) and Nikon Ti2W (Nikon).

The cell distribution was measured from brightfield images of droplets in media on day 0, by mean intensity approximation. The individual droplets were segmented by morphological operations with Canny edge detection and particle analysis (FIJI) and then the mean intensity was measured for each droplet.

Organoid size was measured from brightfield images by segmentation (Supplementary Figure 5a), either by Halo AI for organoids in droplets (Indica Labs, version 3.6) or in Python for organoids in domes. In brief, when using HALO AI, single organoids were automatically detected using a deep learning algorithm (DenseNet AI V2 Plugin) trained to distinguish organoids from background. When using Python, organoids were detected in brightfield images as follows: images were inverted, the scale was normalised by applying quantiles on the image histogram and Multiotsu thresholding. Detected regions were filtered according to size and aspect ratio to filter out segmentation artifacts. All steps in python were performed using algorithm implementations from scikit-image library. Detection was manually validated and those objects which did not correspond to organoids were removed.

Further analysis was performed with Python. The variability in organoid size and cell distribution was expressed in terms of the coefficient of variation, defined as the ratio of the standard deviation divided by the mean value.

### 6. Microfluidic encapsulation

The encapsulation chip was functionalized using 1H, 1H, 2H, 2H-Perfluoro-1-octanol (Merck/Sigma) for 30 minutes, washed with deionized water and dried. The distribution device was primed with 1% bovine serum albumin (BSA) (Miltenyi Biotec) in phosphate buffered saline (PBS) (Gibco) for at least 5 min. Microfluidic encapsulation was performed via flow focusing device as indicated in Supplementary Figure 1a. Pressure and flow rate were controlled with Flow EZ pressure controllers and flow unit (Fluigent).

The 3M™ Novec™ 7500 oil supplemented with 1.5% w/v dSURF surfactant (Fluigent) was injected into the flow focusing device at 15 µL/min. The hydrogel cell suspension was kept on ice and injected at a pressure of 250 or 300 mPa, through a water cooled tube, to the flow focusing device or to the distribution device, respectively. The hydrogel precursor was sheared into monodisperse droplets at the merging points of both fluids in the flow focusing device. The droplets were collected in an Eppendorf tube or a polymerization pool and left at 37 °C for 20 minutes to polymerize.

To remove droplets from oil (Supplementary Figure 1b), the polymerized droplets were passed twice over a hydrophilic PTFE membrane (Merck/Millipore) in a volume of 50-75 µL to enable oil uptake of the PTFE membrane and then droplets were resuspended in desired media, or passed through oil removal device as described below.

### 7. On-chip oil removal

After hydrogel droplets were generated, the droplets were collected in an incubation chamber (Supplementary Figure 7b), placed on top of a heating unit (Inheco), covered with a portable microscope incubator (OkoLab stage top incubator) and polymerized for 20 minutes at 37°C. After polymerization the droplets in oil were introduced into the oil removal device (Supplementary Figure 7c), at the same time 20% (v/v) perfluorooctanol (PFO)–oil solution was pulsated into the device. Both fluids were mixed in the serpentine region of the chip, before entering to the extraction chamber, where a coflow was established between 1% BSA in Hanks’ Balanced Salt Solution (HBSS, Gibco) and the droplet solution, allowing the droplets to transfer to the water based fraction. After oil extraction, the solution containing droplets was collected, spun and replaced with culture media.

### 8. RNA isolation, quantitative PCR and bulk RNA sequencing

For quantitative PCR (qPCR) and bulk RNA sequencing, 2 mL of organoid droplets suspended in media and 2 domes (1:6 diluted, 3.5 µL sized) were collected in 1% BSA-coated Eppendorf tubes, centrifuged and resuspended in a dissociation mixture. The dissociation mixture was composed of 5 mg/mL Collagenase type II (Thermo Fisher Scientific), 1.84 U/mL Dispace (Gibco), 800 U/mL DNase (Merck), 2 µM Y-27632, 0.1% BSA (Miltenyi Biotec) and HBSS (Gibco). The organoid-dissociation mixture was incubated for 30 minutes at 37°C and 500 rpm. Throughout the incubation time, organoids were additionally mechanically dissociated by pipetting every 5 minutes. Dissociated cells were pelleted and resuspended in the desired lysis buffer and frozen at -20°C or -80°C.

For qPCR experiments, 2 mL of organoid droplets suspended in media were used for RNA collection. All material was collected in Eppendorf tubes containing 350 µL RNA lysis buffers (Zymo Research). RNA was extracted using the Quick-RNA™ MiniPrep (Zymo Research) following the manufacturer’s instructions. qPCR analysis was performed using biological duplicates and technical triplicates. cDNA synthesis was performed using the Transcriptor First Strand cDNA Synthesis kit (Roche). qPCR was run on the LightCycler 96 (Roche Diagnostics) using LightCycler® 480 SYBR Green I Master (Roche). β-actin was used as a housekeeping gene. Primers are shown in Supplementary Table 4.

For bulk RNA sequencing, the samples were processed with Organoid DRUG-seq service (ALITHEA genomics). Reads were mapped to the human GRCh38 genome assembly. Differential gene expression analysis was performed using the DESeq2 package^42^.

### 9. Organoid fixation and Histogel embedding

The organoid domes and droplets were gently washed once with 1x PBS and subsequently fixed overnight at 4°C in 4% paraformaldehyde (Sigma). After fixation the material was stored in 4°C, before use. The collected material was embedded into a HistoGel (Epredia) microarray mold, as previously described^43,44^. The molds were processed on an automated tissue processor HistoCore PEARL (Leica) and embedded in paraffin. The formalin-fixed paraffin-embedded (FFPE) blocks were sectioned on a microtome (3 µm) and mounted onto Superfrost® Plus Gold (Epredia).

### 10. Hematoxylin and Eosin Staining

H&E staining was done manually using the H&E Staining Kit (Abcam). The sections were deparaffinized and hydrated in distilled water. The sections were incubated in Hematoxylin for 5 minutes and rinsed two times in water. Then, the sections were incubated in Bluing Reagent for 20 seconds and rinsed and rinsed two times in water. The slides were dipped in absolute alcohol and the excess was blotted off. The section incubated in Eosin Y Solution for 2-3 minutes and rinsed and dehydrated using absolute alcohol. Finally, the slide was cleared, mounted in synthetic resin and scanned by a brightfield whole-slide scanner (Hamamatsu, NanoZoomer S360).

### 11. Multiplex Immunostaining

mIF was automated on Ventana Discovery Ultra automated tissue stainer (Roche Tissue Diagnostics), as described in^43,44^. Slides were baked first at 60°C for 8 minutes and subsequently further heated up to 69°C for 8 minutes for subsequent deparaffinization with Discovery Wash (Ventana). This cycle was repeated twice. Heat-induced antigen retrieval was performed with Tris-EDTA buffer pH 7.8 (CC1, Ventana) at 92°C for 40 minutes. After each blocking step with the Discovery Inhibitor (Ventana) for 16 minutes, the Discovery Inhibitor was neutralized. Primary antibodies (Supplementary Table 5) were diluted in 1X Plus Automation Amplification Diluent (Akoya Biosciences). Primaries were detected using according anti-species secondary antibodies conjugated to HRP (OmniMap anti-Rabbit HRP, Ventana; OmniMap anti-Mouse HRP, Ventana; OmniMap anti-Rat HRP, Ventana; OmniMap anti-Goat HRP, Ventana). Subsequently, the relevant Opal dye (Opal 480, Opal 520, Opal 570, Opal 620, Opal 690, Opal 780, Akoya Biosciences) was applied. After the first sequence, the steps are repeated in each subsequent sequence: Antibody neutralization and HRP denaturation step was applied to remove residual antibody and HRP, Inhibitor, Antibody, Secondary HRP, Opal dye. In the sixth sequence, instead of opal dye, Opal TSA reagent was applied, and immediately after that, Opal dye 780 (Akoya Biosciences) was applied. Lastly, samples were counterstained with 4’,6-Diamidino-2-phenylindole (Ventana). Sequential order of the primary antibodies as well as corresponding dyes were determined during establishment runs. Neutralization of HRP and denaturation of the proteins was performed after every primary antibody cycle in order to avoid cross-bleeding and cross-reacting antibodies. Slides were mounted manually using ProLong™ Gold Antifade Mountant (Thermo Fisher Scientific). Slides were dried for at least 2 hours prior to imaging.

Image analysis of mIF images was performed with HALO AI (Indica Labs, version 3.6). Briefly, single organoids were automatically detected using a deep learning algorithm (DenseNet AI V2 Plugin) trained to distinguish organoids. Following validation,organoids were annotated as individual regions of interest.

The HighPlex FL v.4.2.14 module was used to perform nuclear segmentation based on DAPI+ cells (assisted by HALO’s integrated AI-default ‘nuclear segmentation type’) and specific marker identification. For quantification, DAPI+ nuclei and markers for each distinct cell type of interest were merged (taking membranous and nuclear signals into account). Positive detection thresholds were established for each of the markers taking into account their nuclear or cytoplasm nature and integrated into the HighPlex FL analysis module. The HighPlex FL analysis module was deployed on previously generated regions of interest of the organoids using integration of the classifier in the module. Further analysis was performed with Python.

### 12. Whole mount staining

Permeabilization and blocking were next performed by incubating organoids in 1x PBS containing 0.5% Triton X-100 (Sigma) for 2 hours at room temperature and blocked with 2% normal donkey serum in PBS overnight at 4°C. All stainings were performed in a blocking buffer (2% normal donkey serum in PBS, 0.05% Triton X-100). Samples were incubated with conjugated primary antibody (Supplementary Table 5) for 32-48 hours, washed 3 times for at least 2h with PBS, followed by incubation in blocking buffer containing DAPI (1:1000, Invitrogen), Hoechst (1:1000, Thermo Fisher Scientific) and Alexa Fluor™ Plus 647 Phalloidin (1:2000, Thermo Fisher Scientific) for 24 hours. The organoids were cleared by D fructose (Sigma) for a few hours or overnight and imaged using an CV8000 confocal microscope (Yokogawa). Image contrast was adjusted for visualization purposes.

### 13. Cell viability

Cell viability was assessed by LIVE/DEAD™ Viability/Cytotoxicity Kit (Thermo Fisher Scientific) according to the manufacturer’s instructions. Organoid cultures were washed with DMEM:F12+++ and the Calcein AM, ethidium homodimer-1 and DAPI were added in DMEM:F12+++. The incubation time ranged from 30-90 minutes at 37°C. To stop the staining, a wash with base media was performed. Live/dead staining was visualized on Nikon Ti2 CSU-W1. The maximum intensity z-projection fluorescence images were analysed in FIJI. Positive life threshold and positive dead threshold were established and the number of pixels above the threshold was counted. The viability was expressed as the percentage of life cells where the number of life positive pixels was divided by the total number of positive pixels.

### 14. Drug testing

Tumoroids in droplets, grown in a bioreactor for 12 days, were transferred into a 96-well ultra-low attachment U bottom plate (Primesurface) by a multichannel pipette, and cultured for additional 24 hours in the well plate, before treatment. The tumoroid were treated with 3 individual drugs and one combinatorial drug treatment. The highest drug concentrations were chosen as follows: 1 µM Bortezomib (Tocris), 200 µM SN-38 (Tocris), 50 µM Capecitabine (Selleckchem) and the combinatorial FOLFOX with 50 µM 5-Fluorouracil (Tocris), 2 µM Oxaliplatin (Tocris), 10 µM Leucovorin (Selleckchem). We included a no-treatment control and vehicle control with DMSO. The treatment was performed on technical triplicates and incubated for 48h. Cell viability assessment, imaging and analysis was executed as described above.

### 15. Organoid spotting

Organoids were spotted using the DispenCell device from SEED Bioscience, according to the standard protocol. Shortly, around 400 organoids were resuspended in 1 mL of methylcellulose-based medium, loaded in the machine and dispensed into a 96 ultra low attachment well plate, based on impedance measurements, with the goal to achieve dispersion of 1 organoid per well. A threshold for organoid detection was established manually based on the size of the organoids.

## Acknowledgements

We acknowledge the support of the non profit foundation HTCR, which holds human tissue on trust, making it broadly available for research on an ethical and legal basis. We thank Adrian Filip and Marius Harter for experiment discussions, results interpretation and image analysis support; Max Max Schulz and Ilya Lukonin for support with image analysis. Anke Gehringer for logistic support. We thank Regine Gerard, Ines Silva and Kristina Kromer for help with organoid establishment. We thank Seed Bioscience for support with organoid spotting.

## Author contributions

B.L. and J.L.G.C. conceived the study. The manuscript was written and the data was analyzed by B.L., H.K., and M.C.N with support of R.L.S., J.G.C. and J.L.G.C.. B.L. and H.K. designed and performed most of the experiments with the help of M.C.N and R.L.S. The microfluidic devices were designed, fabricated, and optimized by M.C.N., E.C., and L.B.Z.. I.C. performed mIF stainings.

## Declaration of AI-assisted technologies in the writing process

AI-assisted technologies were used to spell- and language-check written text. Following the use, the text was thoroughly reviewed and revised as necessary, assuming full responsibility for the publication’s content.

## Supplementary Information

**Supplementary Figure 1:**
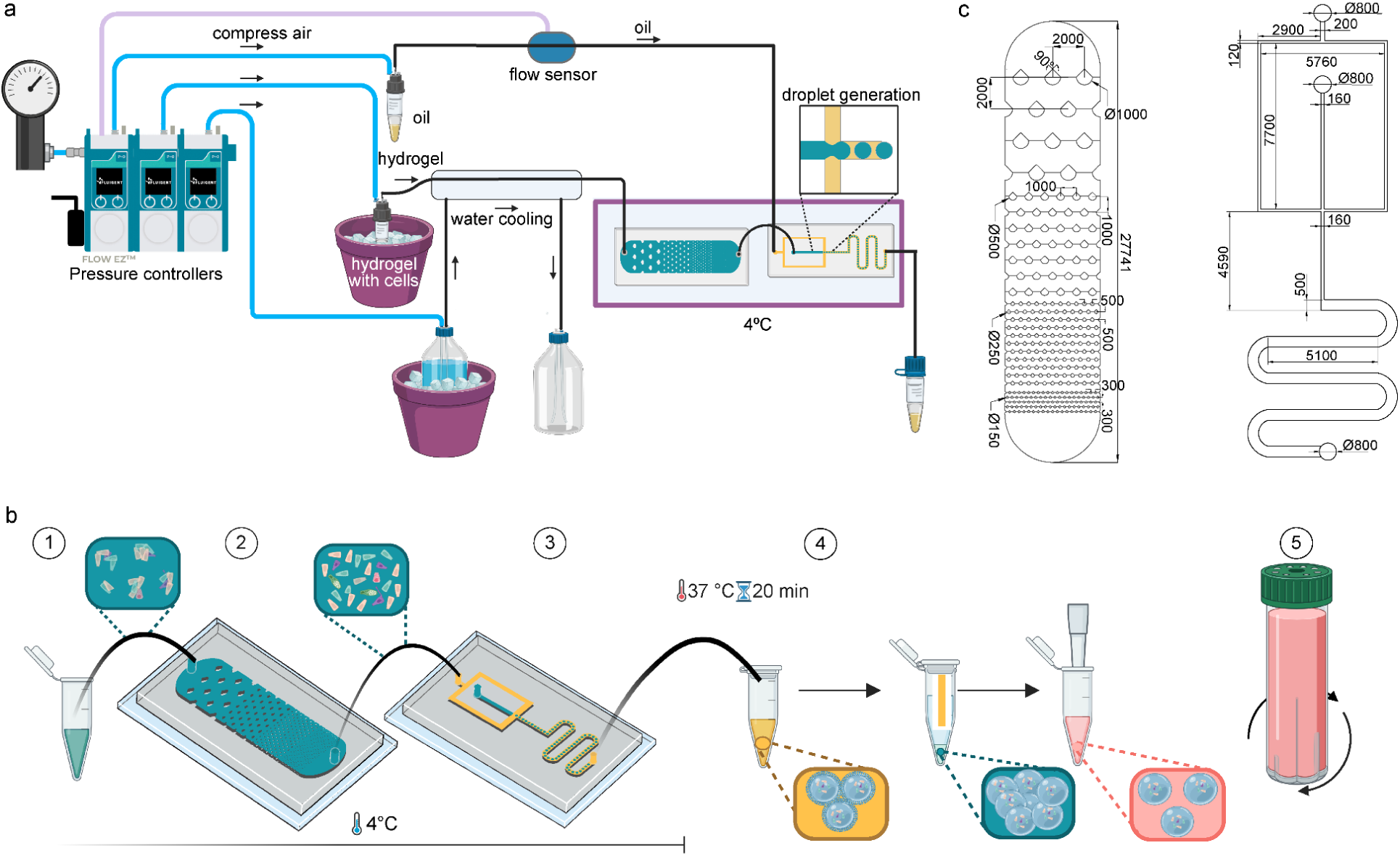
a) Schematic overview of microfluidic set up for droplet generation. b) A schematic of the system workflow. Clumping cell hydrogel suspension (1) is homogenized through the cell distribution device (2) and connected to the flow-focusing encapsulation device (3). Formed hydrogel droplets undergo polymerization, oil extraction and resuspension into media (4). Cell-hydrogel particles are cultured in a bioreactor (5). c) Technical drawings of dissociation chip and flow-focussing device. Schematics were partially created with BioRender.com

**Supplementary Figure 2:**
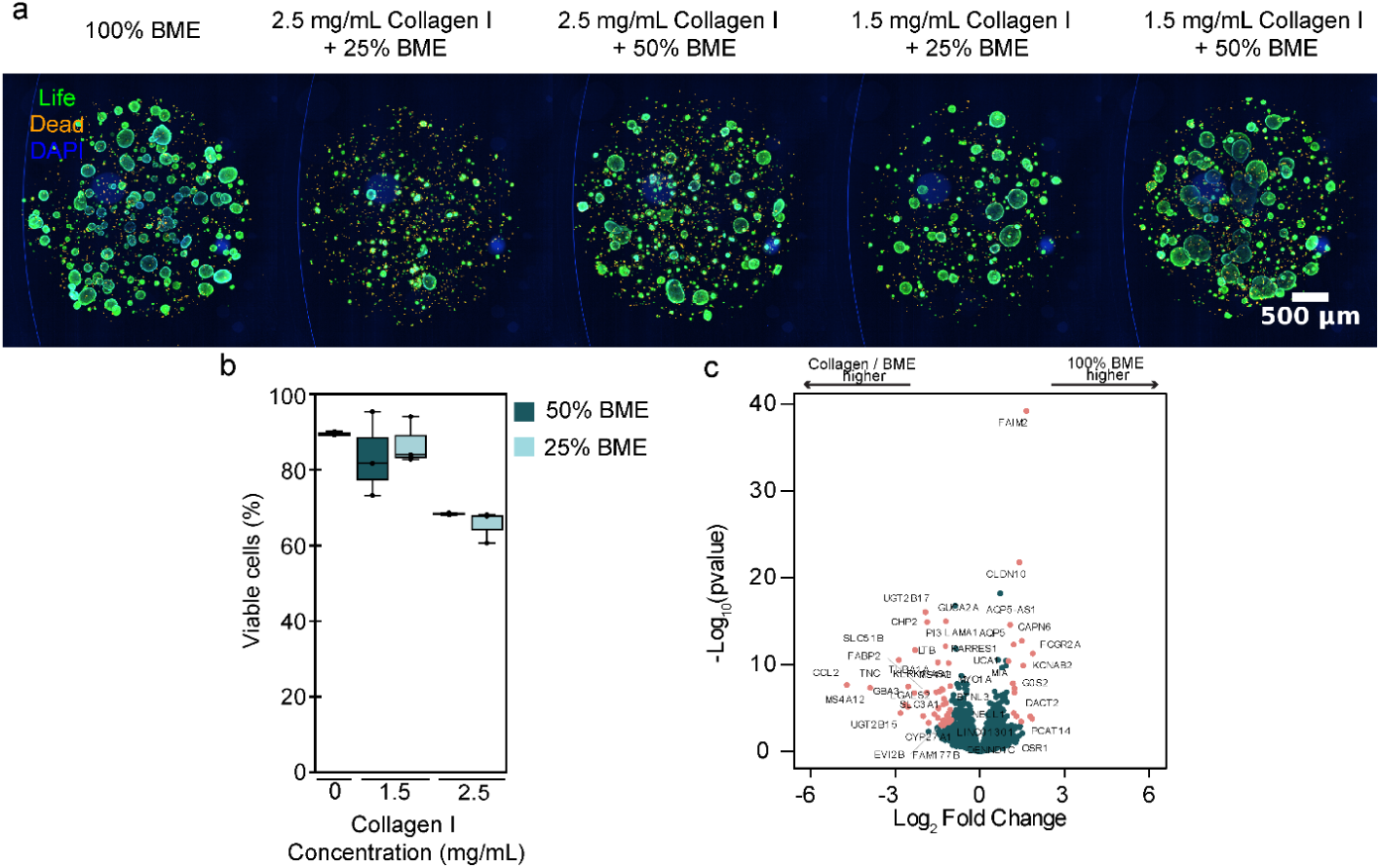
a) Representative fluorescence images of live/dead stainings of organoids growth at different Collagen I / BME (Cultrex) ECM at day 7 (scale bar 500 µm). b) Quantitative fluorescence analysis of cell viability in intestinal organoids at days 7 of culture in domes composed of different ECM by live/dead staining (n_samples_ = 3). c) Volcano plot of differentially expressed genes between 1.5 mg/mL Collagen– 25% BME (Matrigel) ECM mix (left) and BME (Matrigel) (right) subsets (p < 0.05, log2 Fold Change > 1).

**Supplementary Figure 3:**
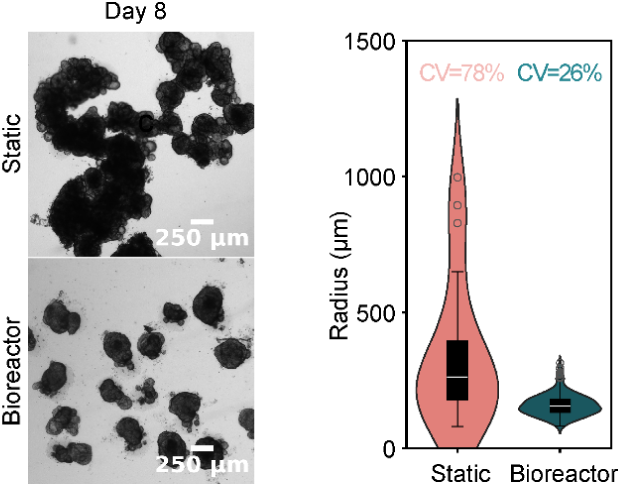
Representative brightfield images of droplets cultured in 6 well plate or bioreactor (Scale bar 250 µm). Size analysis of organoids grown in droplets in static condition or in bioreactor on day 8 with coefficient of variation of organoid size highlighted above (n_static_ = 22, nbioreactor= 237).

**Supplementary Figure 4:**
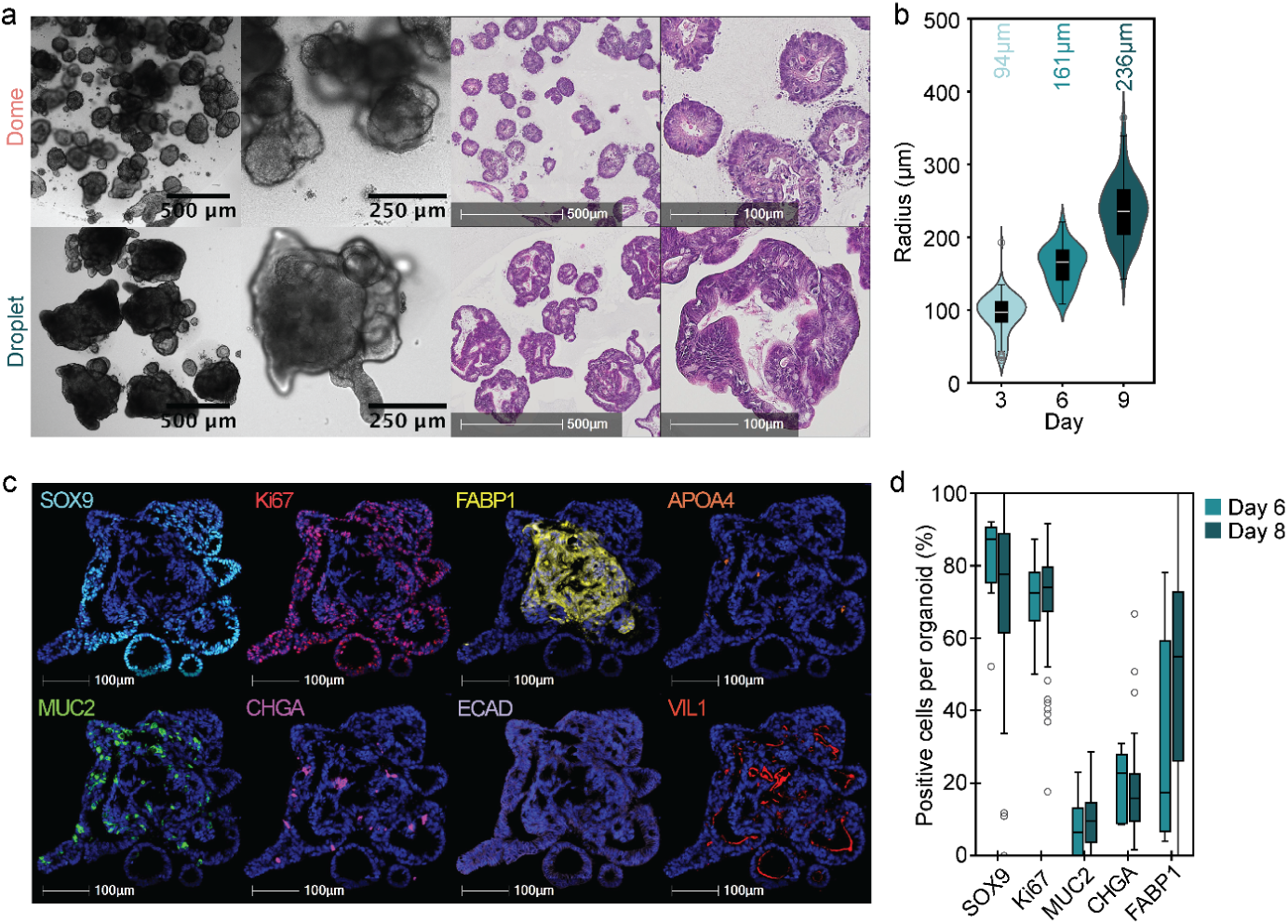
a) Brightfield images and H&E staining comparison of duodenum organoids on day 8 grown in domes or hydrogel droplets. b) Size development analysis of organoids in droplets over several days, with average size highlighted above (n_organoids_ ≥ 55). c) mIF staining of organoids in hydrogel droplets on day 8. d) Quantification of mIF staining of organoids in droplets on day 6 and day 8 (n_organoids_ ≥ 17).

**Supplementary Figure 5:**
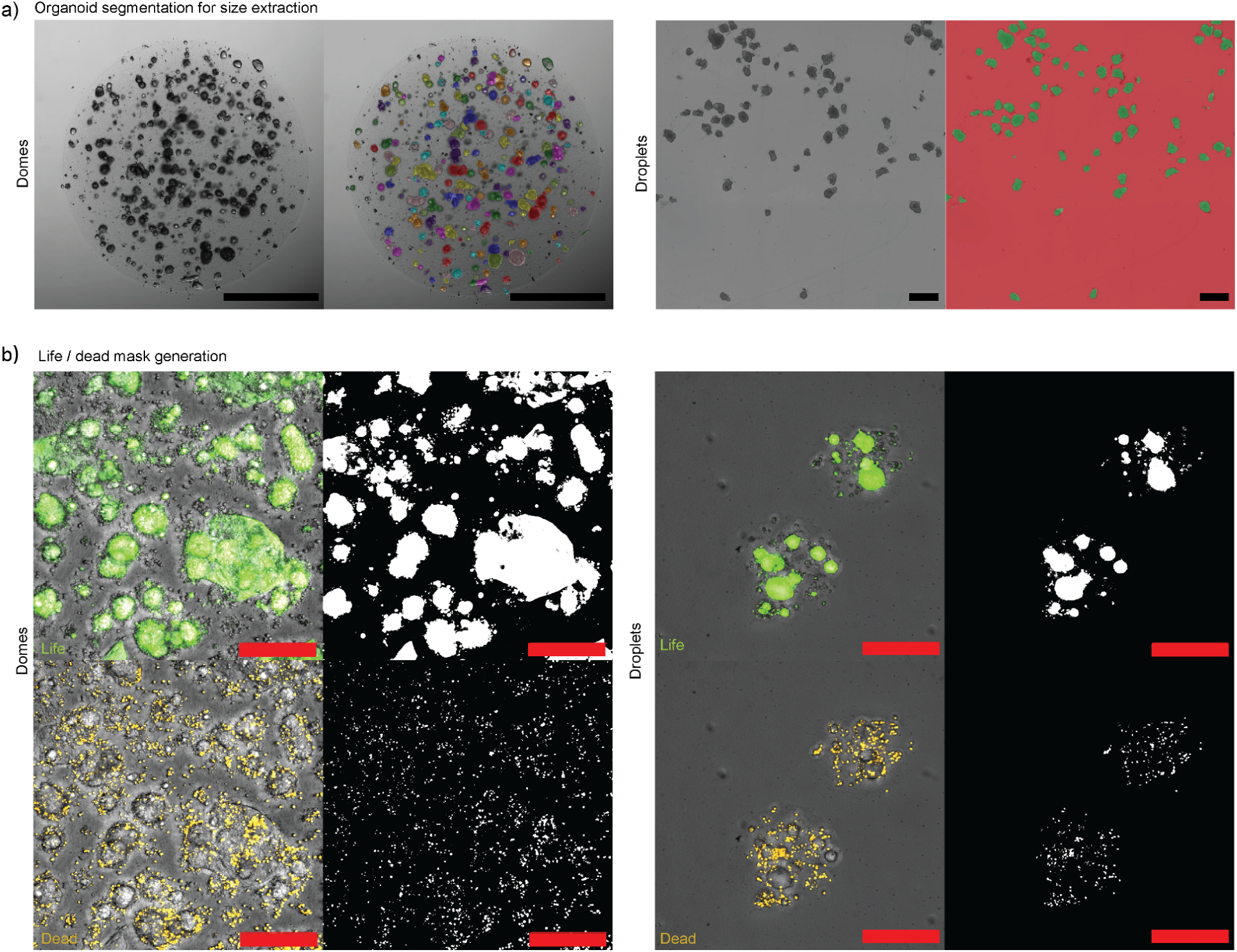
a) Representative images of organoid size analysis. Brightfield image of dome with corresponding color-coded image segmentation masks for individual organoids on left, and brightfield image of organoids within droplets with corresponding green image segmentation masks for individual organoids on right (scale bars 1 mm). b) Representative images of cell viability analysis, overlay brightfield and fluorescence image with the corresponding segmentation mask. Life cells stained by Calcein AM are shown in green and dead cells stained with ethidium homodimer-1 are shown in orange (scale bars 200 µm).

**Supplementary Figure 6:**
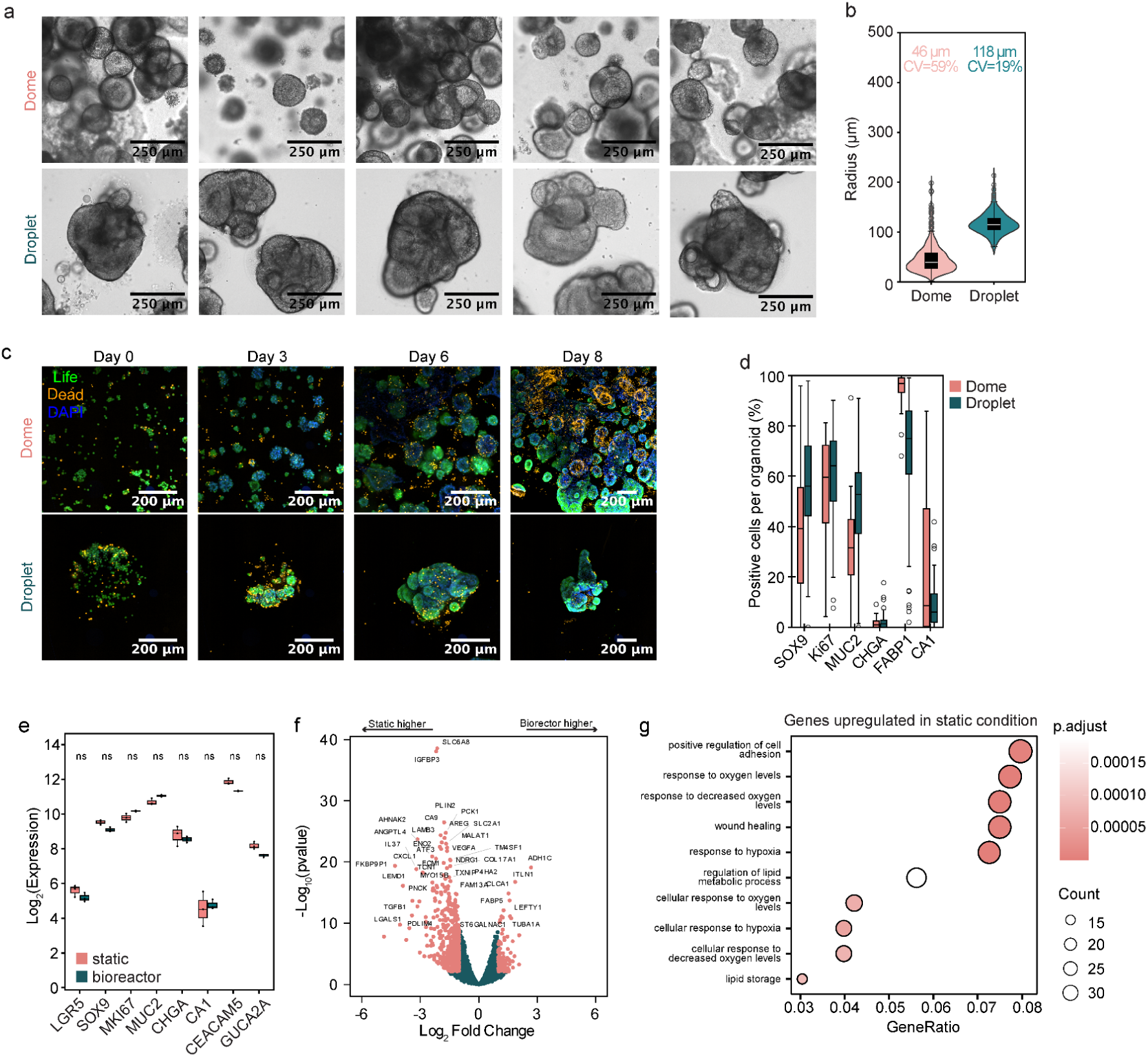
a) Brightfield images of biological replicates of colon organoids on day 8 grown in domes or hydrogel droplets. b) Size analysis of organoids grown in domes or droplets on day 6 with coefficient of variation of organoid size highlighted above (n_organoids_ ≥ 90) c) Representative fluorescence images of live/dead stainings of organoids growth in domes or droplets on day 0, 3, 6, 8 (scale bar 200 µm). d) Quantification of mIF staining of organoids in domes or droplets on day 8 (n_organoids_≥ 67). e-g) Bulk RNA sequencing of organoids grown in droplets in static condition or in bioreactor on day 8 (3 technical replicates). e) Quantification of Log_2_ transformed expression levels. Statistical significance between conditions was assessed using the Wilcoxon test, with p-values adjusted for multiple testing using the Benjamini-Hochberg method. f) Volcano plot of differentially expressed (DE) genes between static (left) and bioreactor (right) condition (p < 0.05, log2 Fold Change > 1). g) Plot showing significantly enriched Gene Ontology biological processes of genes in colon organoids grown in static condition (p-value < 0.05, q-value < 0.2).

**Supplementary Figure 7:**
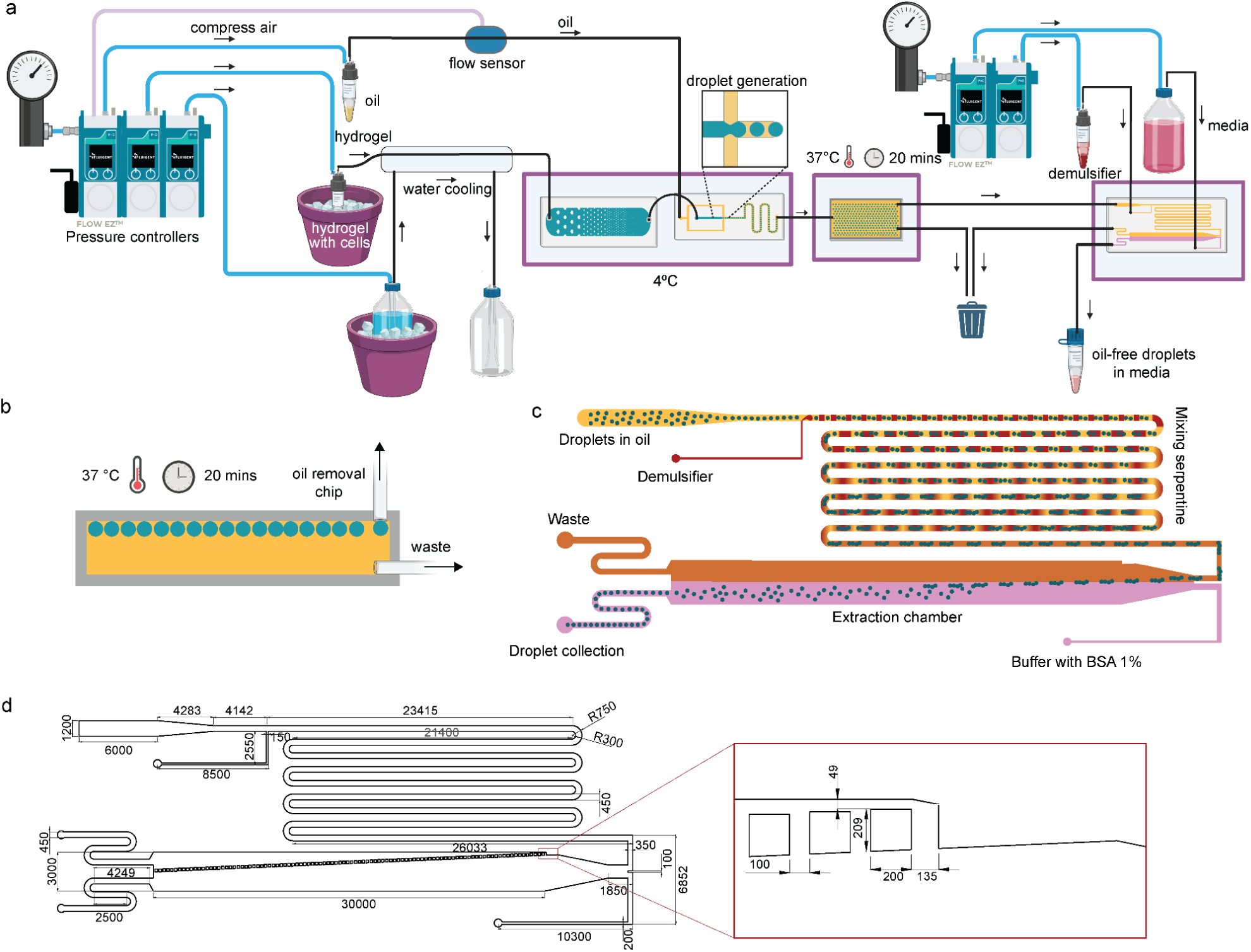
Oil removal microfluidic device: a) Schematic overview of microfluidic set up for droplet generation integrated with polymerization and oil removal module. Schematics were partially created with BioRender.com b) A detailed schematic of the polymerization chamber. c) A detailed schematic of the oil removal chip with workflow description. Schematics were partially created with BioRender.com d) Technical drawings of the flow focusing device.

**Supplementary Figure 8:**
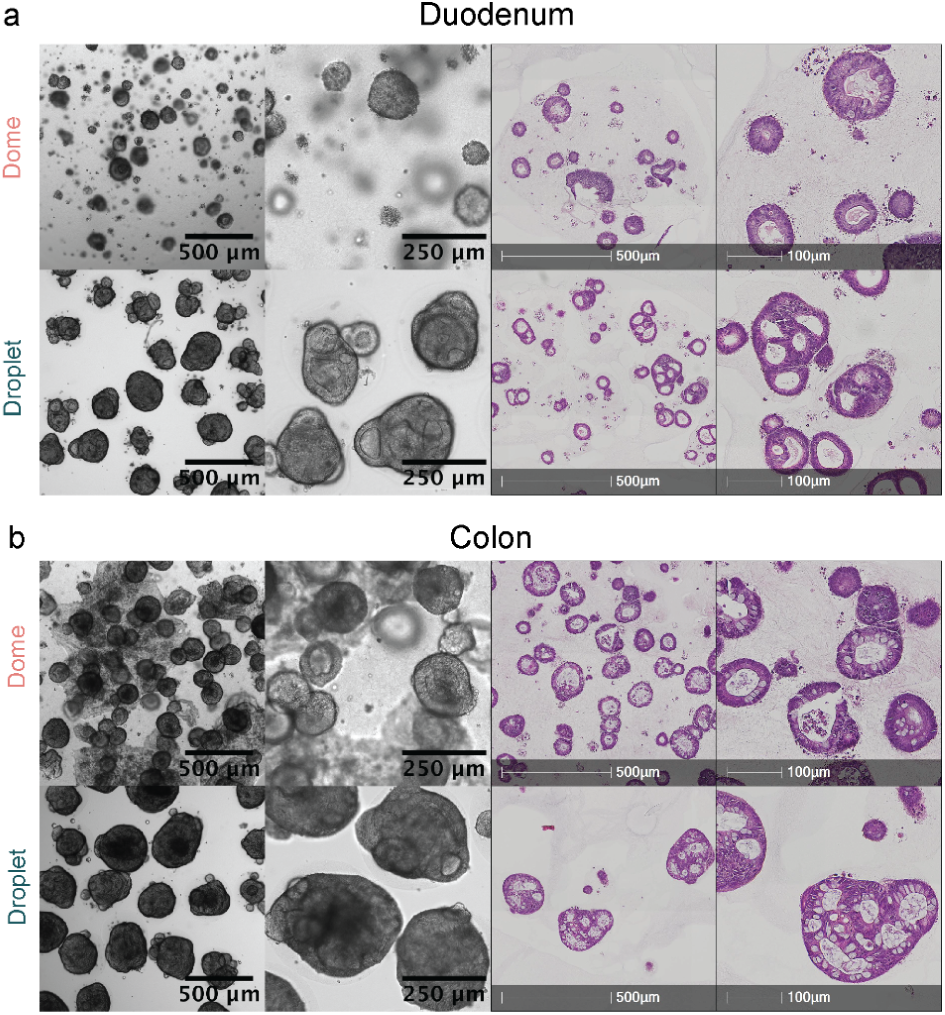
Brightfield images and H&E staining comparison of organoids on day 8 grown in domes or hydrogel droplets generated with the oil removal device: a) organoids derived from duodenum, b) organoids derived from colon (scale bars as indicated 500 µm, 250 µm, 100 µm).

**Supplementary table 1:** Material list.

| Material | Catalog Number NUMBER | Supplier |
| --- | --- | --- |
| SU-8 Photoresist | GM1075 | Gersteltec Sàrl |
| Propylene Glycol Monomethyl Ether Acetate | 484431 | Sigma Aldrich |
| 1H, 1H, 2H, 2H-Perfluoro-1-octanol (PFO) | 370533-25G | Merck/Sigma |
| Chlorotrimethylsilane | 92361-250ML | Sigma |
| MycoAlert™ PLUS Mycoplasma Detection Kit | LT07-705 | Lonza |
| Advanced DMEM:F12 | 12634-010 | Gibco |
| HEPES | 15630-056 | Gibco |
| Glutamax | 35050-038 | Gibco |
| Penicillin-Streptomycin | 15140-122 | Gibco |
| Gentle Cell Dissociation Reagent | 100-0485 | Stemcell Technologies |
| Tissue Culture Treated Well Plates | CC7682-7524 | CytoOne® |
| COSTAR Plastic Ultra-Low 6-well plate | 3471 | Corning |
| Growth Factor Reduced Matrigel | 356231 | Corning |
| TrypLE™ Express Enzyme | 12604-013 | Gibco |
| DNAse | 4536282001 | Roche |
| Y-27632 | 72305 | STEMCELL Technologies |
| DL-Dithiothreitol Solution (DTT) | 43816-10mL | Sigma-Aldrich |
| 40 µM Cell Strainer | 43-50040-01 | Pluriselect |
| Biomatrix TelCol-6 (Collagen I) | 5225-50mL | Biomatrix |
| Neutralization Solution | 5229-10mL | Biomatrix |
| Hydrophilic PTFE Membrane | BGCM10448213 | Merck/Millipore |
| IntestiCult™ Organoid Growth Medium (Human) | 6010 | StemCell Tech |
| Primocin | ant-pm-1 | Invivogen |
| Collagenase type II | 17101015 | ThermoFisher |
| Dispase | 17105-041 | Gibco |
| MACS® BSA Stock Solution | 130-091-376 | Miltenyi Biotec |
| HBSS | 14175-095 | Gibco |
| RNA Lysis Buffer | R1060-1-100 | Zymo Research |
| Quick-RNA™ MiniPrep | R1055 | Zymo Research |
| Transcriptor First Strand cDNA Synthesis Kit | 4897030001 | Roche |
| LightCycler® 480 SYBR Green I Master | 4707516001 | Roche |
| HistoGel | HG-4000-012 | Epredia |
| Hematoxylin and Eosin Staining Kit | ab245880 | Abcam |
| Discovery Wash | 950-510 | Ventana |
| Tris-EDTA Buffer | 950-227 | Ventana |
| Discovery Inhibitor | 760-4840 | Ventana |
| OmniMap anti-Rabbit HRP | 760-4311 | Ventana |
| OmniMap anti-Mouse HRP | 760-4310 | Ventana |
| OmniMap anti-Rat HRP | 760-4457 | Ventana |
| OmniMap anti-Goat HRP | 760-4647 | Ventana |
| Opal 480 | FP1500001KT | Akoya Biosciences |
| Opal 520 | FP1487001KT | Akoya Biosciences |
| Opal 570 | FP1488001KT | Akoya Biosciences |
| Opal 620 | FP1495001KT | Akoya Biosciences |
| Opal 690 | FP1497001KT | Akoya Biosciences |
| Opal 780 | FP1501001KT | Akoya Biosciences |
| DAPI | 760-4196 | Ventana |
| ProLong™ Gold Antifade Mountant | P36930 | Thermo Fisher Scientific |
| LIVE/DEAD™ Viability/Cytotoxicity Kit | L3224 | Thermo Fisher Scientific |
| Alexa Fluor™ Plus 647 Phalloidin | A30107 | Thermo Fisher Scientific |
| 3M™ Novec™ 7500 oil | HFE-7500 3M | Fluigent |
| dSURF surfactant | ODSURF5G | Fluigent |
| Donkey serum | S30-M | Sigma |
| 5-Fluorouracil | 3257 | Tocris |
| Oxaliplatin | 2623 | Tocris |
| Leucovorin | S1236 | Selleckchem |
| Bortezomib | 7282 | Tocris |
| SN-38 | 2684 | Tocris |
| Capecitabine | S1156 | Selleckchem |
| DAPI | 62248 | Thermo Fisher Scientific |
| Hoechst | 62249 | Thermo Fisher Scientific |
| DMSO | D2650 | Sigma-Aldrich |

**Supplementary table 2:** Donor information for Intestinal resection.

| Donor number | Anatomical origin | Sex | Age | Biopsy origin |
| --- | --- | --- | --- | --- |
| AN96289 | Healthy colon ascendens | female | 50 | HTCR |
| AN77238 | Healthy duodenum | female | 61 | HTCR |
| P756 | Healthy colon ascendens | male | 78 | University Hospital Basel |
| P673 | Healthy Colon Cecum | male | 75 | University Hospital Basel |
| P599 | Colon Adenocarcinoma | male | 55 | University Hospital Basel |

**Supplementary table 3:** Composition of intestinal organoid media.

| Components | Vendor | Reference | Concentration |
| --- | --- | --- | --- |
| Advance DMEM/F12 | Thermo Fisher Scientific | 12634010 |  |
| Glutamax | Thermo Fisher Scientific | 35050-061 | 2 mM |
| Penicillin-Streptomycin | Thermo Fisher Scientific | 15140122 | 1x |
| HEPES | Thermo Fisher Scientific | 15630080 | 10 mM |
| Rspo3-CM | ImmunoPrecise Antibodies | R001 | 2% |
| B27 min VitA | Thermo Fisher Scientific | 17504044 | 1x |
| Noggin-CM | Immunoprecise antibodies | N002 | 2% |
| N-acetyl-L-Cysteine | Sigma | A9165-5G | 1mM |
| Primocin | Invivogen | ant-pm-1 | 100 µg/mL |
| IGF-1 | BioLegend | 590908 | 100 ng/mL |
| FGF2/FGF-basic | R&D | 3718-FB | 50 ng/mL |
| EGF | Peprotech | AF-100-15 | 50 ng/mL |
| A83-01 (ALK4/5/7 inhibitor) | Tocris | 2939 | 500 nM |
| Wnt surrogate | ImmunoPrecise Antibodies | N001 | 0.15 nM |
| Gastrin | MedChem Express | HY-P1097 | 10 nM |
| Y-27632 dihydrochloride (Rho-k inhibitor) | STEMCELL Technologies | 72305 | 10 µM |

**Supplementary table 4:**
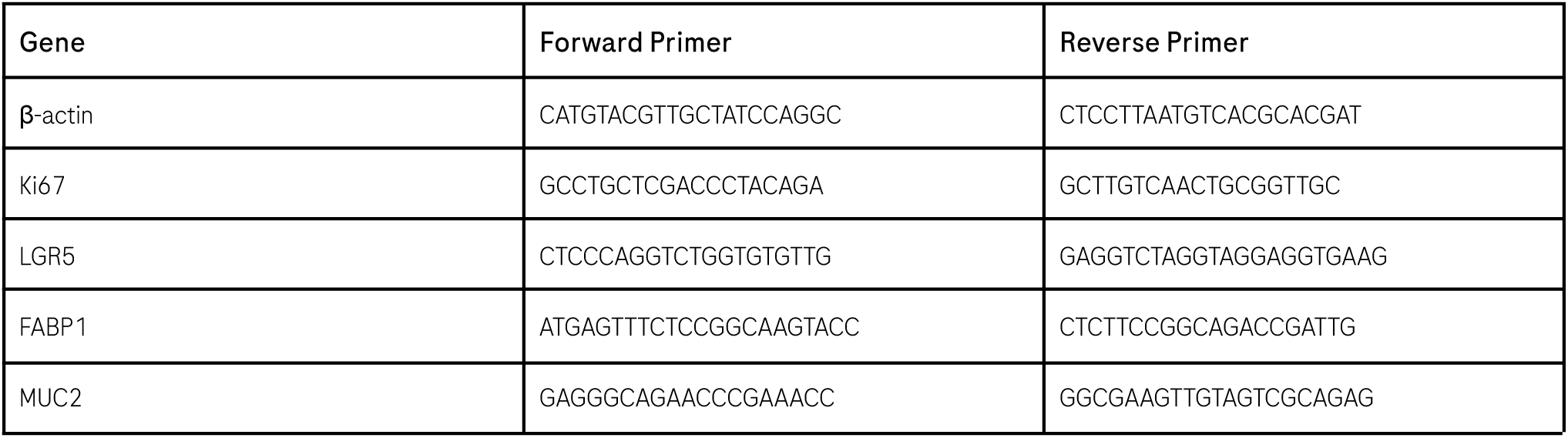
Forward and reverse primer used for qPCR.

**Supplementary table 5:**
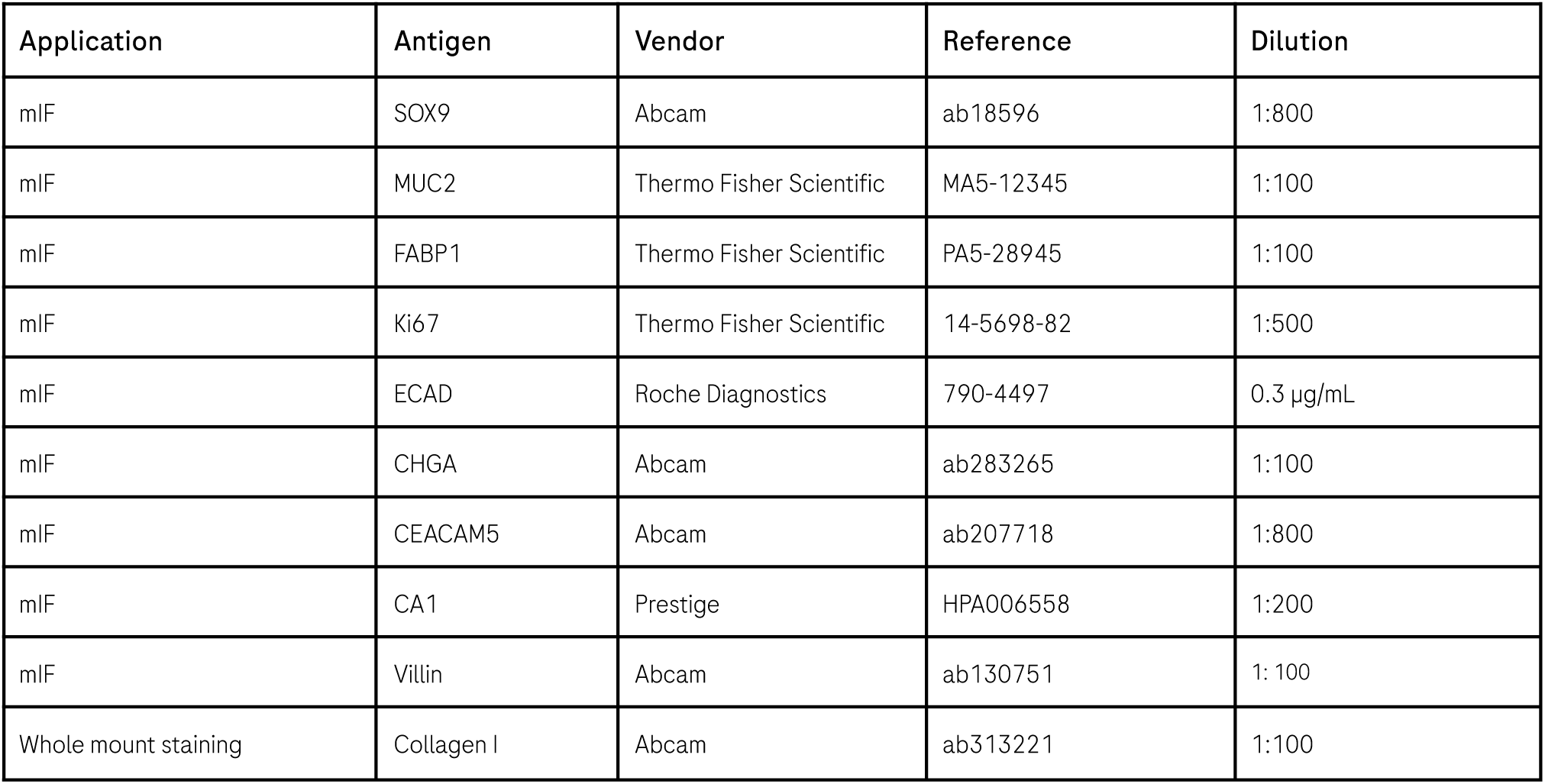
List of antibodies used for Multiplex Immunofluorescence (mIF) antibodies and whole mount stainings.

